# A link between genotype and cellular architecture in microbiome members as revealed by cryo-EM

**DOI:** 10.1101/2022.09.08.507075

**Authors:** Benedikt H Wimmer, Sarah Moraïs, Ran Zalk, Itzhak Mizrahi, Ohad Medalia

## Abstract

Microbial taxonomy is not yet sufficient to describe microbe functionality and ecology. Since function is often linked to structure, we sought here to use cryo-electron microscopy and tomography to analyze microbial cellular architecture and correlate it to specific phylogenies and genomes. We cultured and imaged a large collection of microbiota covering 90% of the richness of the core rumen microbiome at the family level, which we selected as a model for our analyses. Based on measurements of several parameters, we found that the structural similarity of microbiota is significantly related to their taxonomic distance, i.e., closely related microbes have similar cellular architectures. However, above the *Family* level, these similarities end: the structural diversity stops increasing with phylogenetic distance. Our results highlight that cellular architectures could serve as an important parameter in microbial ecology and microbial ecosystems.

## Introduction

There is an enormous discrepancy between microbial taxonomy and functionality ^1,2^. Bacterial species have been defined empirically as clusters of similar organisms, combining both phenotype and common metabolic capabilities ^3^. Microbial systematics were later modified to combine phenotypic, genotypic, and sequence-based phylogenetic data within a framework of standards and guidelines for describing and identifying prokaryotes ^4^, but the definition of prokaryotic species remains controversial ^5,6^. Deepening the link between taxonomy and functionality is essential for the studies of microbial ecology and ecosystems to better interpret and scrutinize the microbial features that govern community composition and dynamics.

Most current workflows for microbiome studies are based on the analysis of phylogenetic composition as indicated by 16S rRNA sequencing, frequently complemented by metagenomic sequencing, to link functionality and phylogeny. The conventional morphological classification used to describe microorganisms is still at the level of the one used by Antoni van Leeuwenhoek in 1676 using a very simplistic light microscope ^7^. This description classifies all phylogenies into a few discrete shapes such as rods, spheres and spirals. These visual phenotypes are surely not sufficient to reflect the uniqueness of different phylogenies and are mostly limited by the low resolution of conventional brightfield light microscopy.

Thus, we explored if the cellular architecture of microorganisms directly determined by high-resolution electron microscopy can expand our comprehension of microbial phylogeny, physiology and ecology within an ecosystem. Cryo-EM allows the direct inspection of unstained biological materials in a near-native state at nanometer resolutions. Very thin biological samples, for example purified proteins or viruses, are applied to a sample carrier and rapidly cryo-fixated. Through repeat observations of randomly distributed particles within the sample and alignment and averaging of those observations, atomic-resolution 3D structures can be determined ^10^.

Cryo-electron tomography (cryo-ET) allows to retrieve the three-dimensional (3D) structure of individual heterogenous objects of interest such as cells ^11,12^. Projections of an object at multiple tilt angles are acquired and back-projected *in silico* to reconstruct its 3D architecture ^13,14^. Cryo-ET has opened a new window into the intra-and extracellular organization of prokaryotic cells, in some small bacteria even enabling a complete survey of the cells’ contents ^15^. It was used to uncover the architecture of chemoreceptor arrays ^16^, contractile injections systems ^17^, surface layers ^18^ and proteins supporting biofilm formation *in situ* in a close-to native state ^19^. While cryo-ET has been instrumental in understanding the most important assemblies and features of prokaryotic cells, few studies have employed it to elucidate their emergence and evolution. Such studies highlighted the potential of directly observing the outcomes of evolutionary processes through cryo-ET, e.g. Chaban and colleagues compared subtomogram averages of flagellar motors in proteobacteria and traced the higher torque to an increased number of stator complexes in epsilon-proteobacteria ^20^. In an elaborated study, Kaplan *et al*. surveyed tens of thousands of tomograms of ∼90 species to create a taxonomy of outer membrane projections in diderm bacteria^21^. Recently, a team at Institut Pasteur combined genomics research with cryo-ET to reconstitute the emergence of monoderm bacteria through the loss of membrane tethers ^22^.

Here, we analyze 69 representative strains of gut microbiota by means of cryo-EM and -ET to evaluate our hypothesis of a link between cellular architecture, phylogeny and ecology. Using a set of five structural parameters, we demonstrate that bacterial morphology is responsive to changes in the environment such as the type of carbon source. Furthermore, by comparing structural similarity across taxonomic relationships, we show that the cellular architecture is relatively conserved within a genus, but not in higher taxonomic levels. In this context, we closely inspect members of the *Lachnospiraceae* family, which cluster together by genome contents, but not by cellular architecture. We hypothesize that this divergence between phylogeny, genotype and morphology reflects ecological constraints, leading to specialized cellular architectures within a family or parallel emergence of similar features in unrelated phylogenies. Therefore, the cellular architecture represents an important parameter that could help better understand microbial phylogeny, microbe-environment interactions in microbial ecology and microbial ecosystems.

## Results

### The core rumen microbiome as a model for large scale microbial structural characterization

The microbial population of the rumen exhibits a plethora of cellular morphologies. To make this diversity accessible to future studies of microbial ecosystems, we cataloged it by cryo-EM and - ET. The core rumen microbiome serves as a model to correlate structural microbial characteristics of a full ecosystem to phylogeny and genomics. We gathered a large microbial isolate collection covering 90% of the core rumen microbiome diversity at the order level (microorganisms present in 80% of a 1000 cow cohort ^23^). Our collection enriches this taxonomic diversity with phylogenies generally encountered in other gut microbiomes, such as Actinobacteria, Proteobacteria, Fusobacteria and Campylobacteriota. We cultured 69 microbial species spanning over 45 genera and 10 phyla (Fig. 1 and Supplementary Fig. 1). In addition, we performed new whole-genome sequencing on 10 of the cultured strains where whole-genome data was not available.

**Figure 1:**
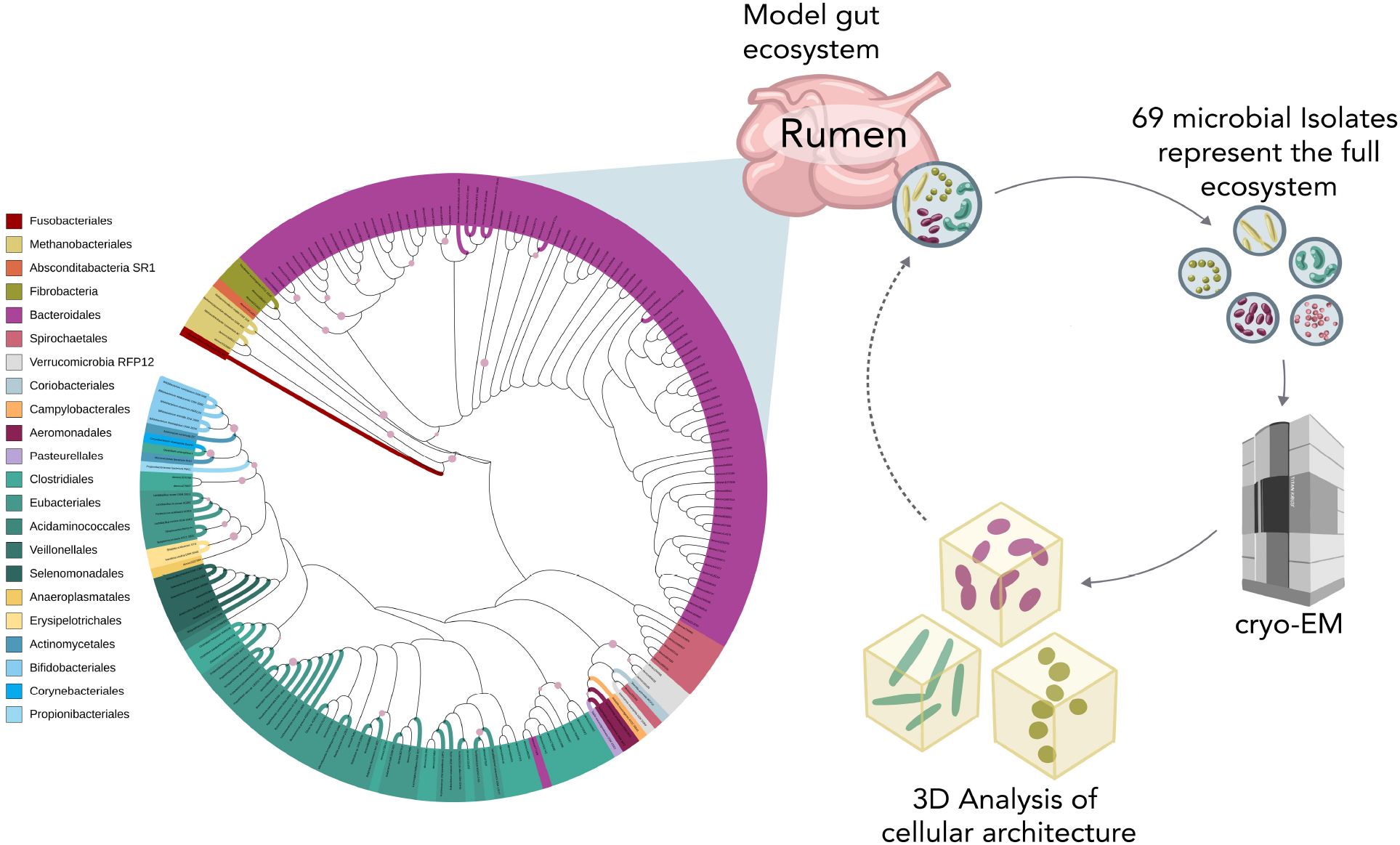
A diverse ecosystem studied by cryo-ET. To study the link between ecology, taxonomy and cellular architecture, the rumen core microbiome was selected as a model. A selection of 69 isolates (colored branches in the phylogenetic tree) was subjected to cryo-electron microscopy and subsequent three-dimensional analysis of cellular architecture (subset of 40 suitable isolates). This analysis yields insights into the structural composition of the model gut ecosystem. The phylogenetic tree is based on the 16S rRNA gene of core microorganisms present in 80% of the individuals at the order level in a cohort of 1000 cows. Color coding is according to order level. Bootstraps with a confidence higher than 95% are displayed as pink circles. The tree was created using iTOL ^49^.

To probe the relationship between ecology and structural characteristics, we challenged some microbes by culturing them on several carbon sources, resulting in 75 samples overall. Sample thickness is a major limitation in cryo-EM, therefore not all 69 strains could be analyzed to the same degree. Here, we acquired and analyzed cryo-ET data of 38 bacterial and two archeal strains. For the remaining 29 strains, only single projection images could be acquired and therefore some structural parameters could not be resolved (Supp. Tab. 1). However, analysis levels do not cluster with taxonomy yielding a good coverage of all phylogenies in the high-resolution dataset (Supp. Fig. 1).

### The core rumen microbiome spans a wide range of cell shapes and sizes

To gain an initial overview of the morphological diversity within the collection, we initially acquired ca. 4,500 electron micrographs at varied magnifications of 3,600x - 11,500x (examples in Fig. 2, Supp. Fig. 2). At these magnification levels, microbes could easily be distinguished by their characteristic shape and dimensions.

**Figure 2:**
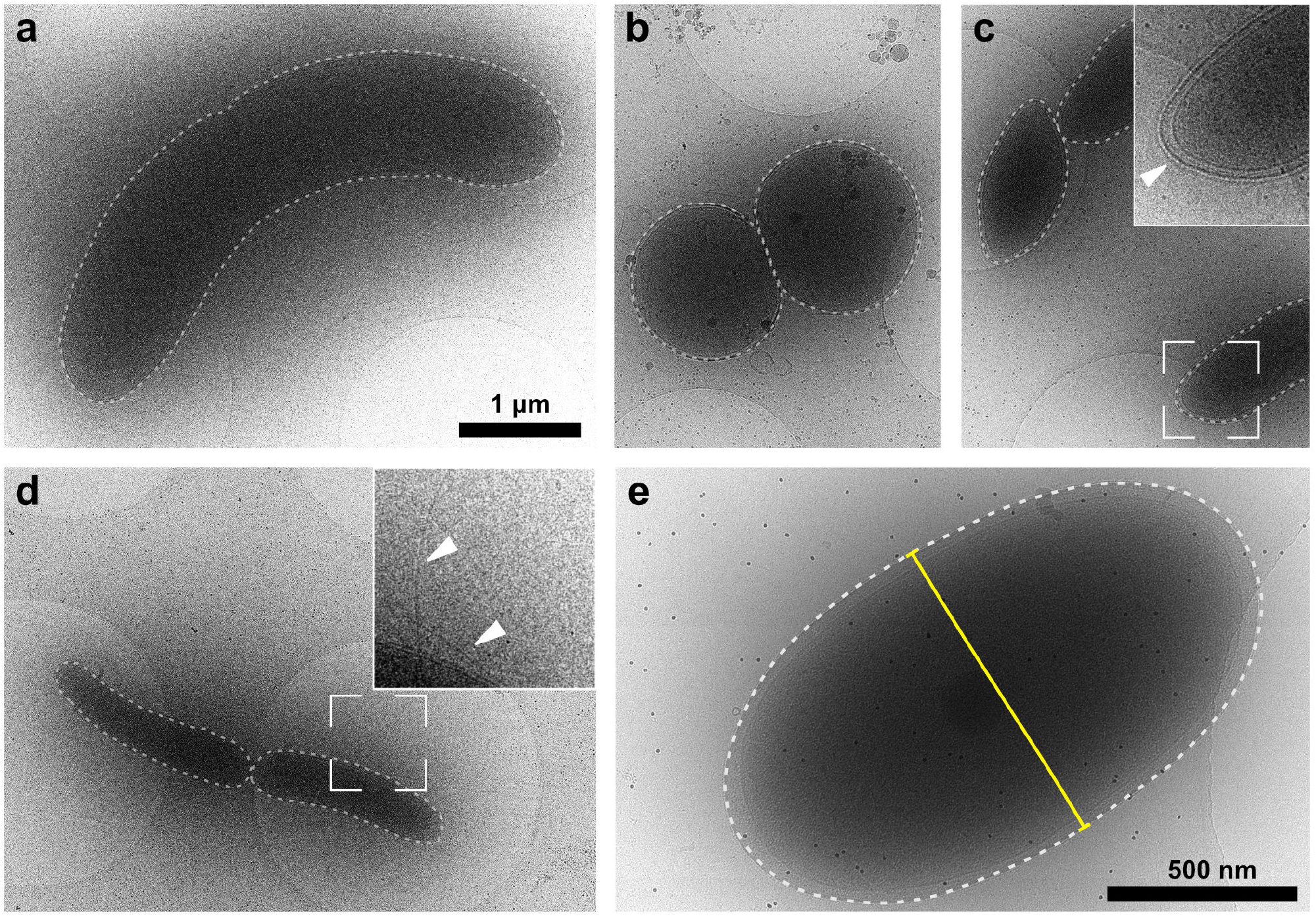
Bacteria of the core microbiome span a plethora of morphologies. Selection of representative micrographs of microbiome members. (a) *Selenomonas ruminantium* AB3002, a diderm rod-shaped bacterium isolated from the bovine rumen. (b) *Succinimonas amylolytica* DSM 2873, a diderm coccoid bacterium isolated from the bovine rumen. (c) *Prevotella ruminicola* ATCC 19189, a diderm bacterium isolated from the bovine rumen. Here, an S-Layer is visible in the micrographs (insert, arrow). (d) *Agathobacter ruminis* DSM 29029, a monoderm vibrioid bacterium isolated from sheep rumen. Flagella are visible in the micrographs (insert, arrows). (e) High-exposure micrograph of *Akkermansia muciniphila* DSM 22959, an important player in the human intestinal microbiome linked to metabolic benefits in the early developing cow rumen ^50,51^. To compare shapes, the outline was manually traced (dotted line) and the circularity of the shape calculated in FIJI. Furthermore, the diameter was measured (yellow line). Panels (a) - (d) at the same magnification.

A total of 45 microbes (65% of the analyzed strains) were classified as rod-shaped (eg. *Selenomonas ruminantium* AB3002 or *Akkermansia muciniphila* DSM 22959, Fig. 2a/e), 10 microbes (14%) as ovoid (eg. *Prevotella ruminicola* ATCC 19189, Fig. 2c), 8 microbes (12%) as coccoid (eg. *Succinimonas amylolytica* DSM 2873, Fig. 2b) and 6 microbes (9%) as vibrioid (eg. *Agathobacter ruminis* DSM 29029, Fig. 2d). For some samples, further features were observed at this magnification, such as S-Layers (Fig. 2c, insert) or flagella (Fig. 2d, insert).

To compare the cellular architecture across species, the diameter and shape were quantified as outlined in Fig. 2e (full results in Supp. Table 1). The diameter was measured at the center of the individual cell, orthogonal to the long axis. The average diameter of the strains in the screening collection was 0.8 µm (range 0.38 - 1.81 µm). The shape was quantified by tracing the cell outline manually, fitting the points with a spline curve and then calculating the circularity, which correlated highly with manual shape classification (One-way ANOVA, F = 58.39, p < 0.0001).

### Phylogenies can be distinguished using five measures of cellular architecture

To better analyze the cellular architecture, we applied cryo-ET to all samples with suitably thickness, i.e. < 600 nm in edge regions. Obvious features were manually annotated using reference images from the Cell Structure Atlas online textbook ^24^. Tomograms from *Pseudobutyrivibrio sp*. LB 2011 and *Wolinella sp*. ATCC 33567 are shown in Fig. 3 (representative slices from all samples in Supplementary Fig. 2).

**Figure 3:**
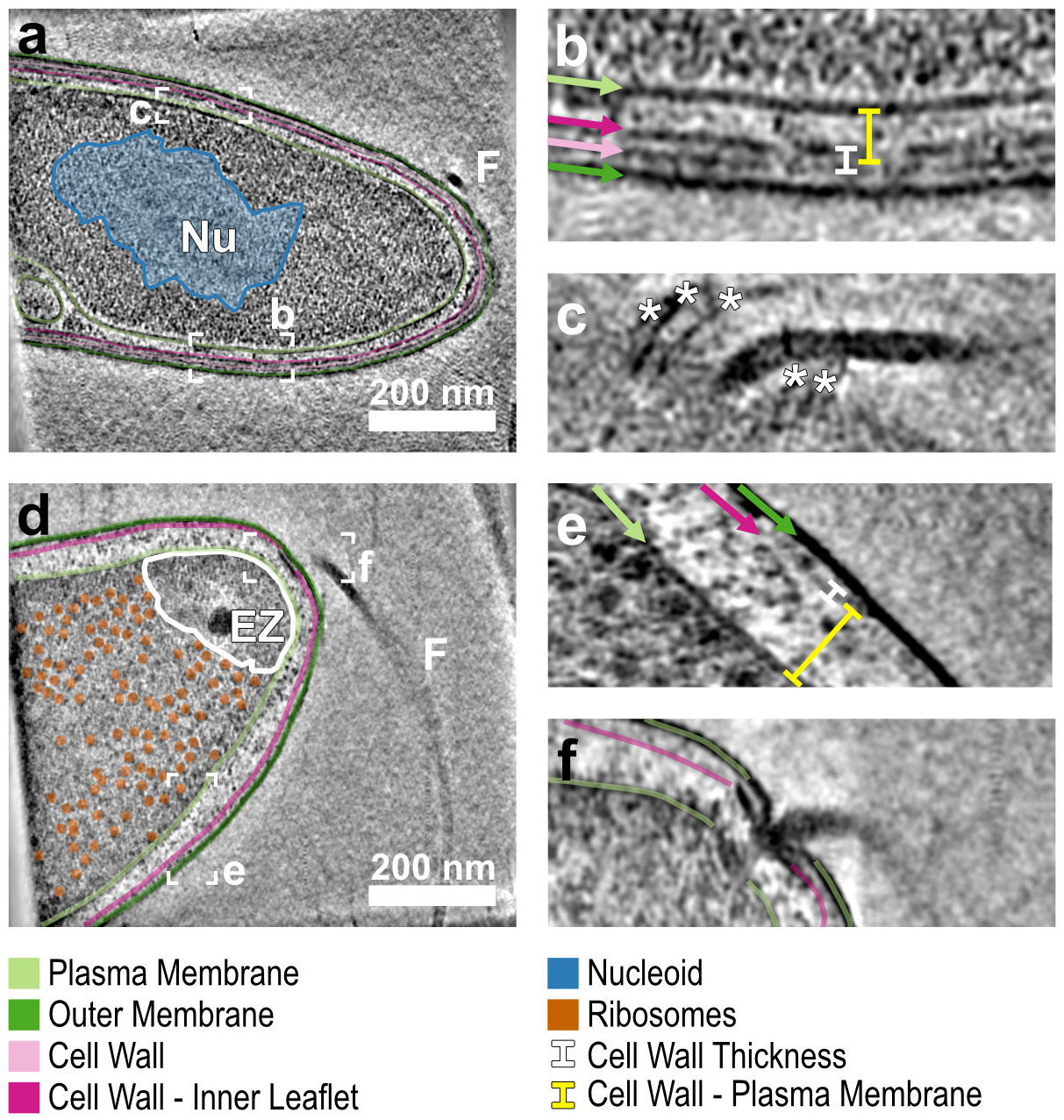
cryo-ET elucidates the cellular architecture. Central cross-sections through tomograms of bacteria reveal the cellular architecture. (a) *Pseudobutyrivibrio sp*. LB 2011 contains a dense cytoplasm, the nucleoid DNA stands out by its stringy texture (blue overlay, Nu). Plasma membrane (PM, light green), cell wall (CW, two leaflets in pink) and outer membrane (OM, dark green) are highlighted. A flagellum (F) is visible next to the cell. (b) Closer inspection of the cell surface reveals a diderm bacterium with a cell wall in two leaflets (pink), membranes are highlighted in light green (PM) and dark green (OM). The cell wall thickness (white line) and distance from the plasma membrane (yellow line) were measured as indicated. (c) Tracing the flagellum leads to a top view of the flagellar motor adjacent to several pili (stars). (d) A central cross-section through a *Wolinella sp*. ATCC 33567 cell. Again, PM, CW and OM are traced. Ribosomes are visible throughout the cytoplasm (orange), with the exception of a polar exclusion zone (white outline, EZ). A flagellum emerging from the tip is visible next to the cell (F). (e) PM and OM are clearly visible bordering an electron-light periplasm. The CW is visible adjacent to the OM as a faint band. Measurements were performed as indicated. (f) The flagellar motor is clearly visible in cross-section. Slice thickness 1 nm.

In a central x-y section of the tomogram (Fig. 3a), a dense cytoplasm enclosed by the plasma membrane (PM, light green) could easily be distinguished from the lighter periplasm framed by the outer membrane (OM, dark green). The PM, the cell wall (two leaflets, pink) and the OM could be traced along the entire field of view. In some samples, the nucleoid (blue overlay) stood out from the cytoplasm through its filamentous texture. The cell surface organization was inspected carefully. In the case of the *Pseudobutyrivibrio sp*., three continuous layers were observed outside the PM (Fig. 3b, light green): a 13.4 ± 0.8 nm thick cell wall consisting of two parallel layers (pink) at a distance of 14.6 ± 0.6 nm and an OM (dark green) 28.8 ± 1.5 nm removed from the PM. Flagella and pili were also frequently observed, here as a top view (Fig. 3c).

For *Wolinella sp*., cyto- and periplasm again are easily distinguishable (Fig. 3d). In the cytoplasm, ribosomes are distributed throughout (orange), with the exception of a polar exclusion zone (EZ, white outline), which contains the flagellar motor (Fig. 3f) and a top view of a chemoreceptor array (not shown). The periplasm itself is much less dense than that of *Pseudobutyrivibrio sp*., and contains only one very light continuous cell wall layer close to the OM (Fig. 3d). The flagellar motor is clearly visible in cross-section (Fig. 3f). The stator scaffold spanning the periplasmic space initially described by Chaban *et al*. ^*20*^ is clearly visible, as are the anchors in the PM, OM and cell wall.

Even though macromolecular complexes such as flagella, chemoreceptor arrays, ribosomes and nucleoids were frequently observed in the tomograms, we decided against parametrization based on the presence or absence of specific complexes as only a part of the cell volume could be surveyed using tomography. Thus, we determined three parameters related to the surface structure of the cells: the presence or absence of an outer membrane (OM), the cell wall thickness and distance of the cell wall from the plasma membrane.

To quantify the similarity of cellular architectures between samples, we calculated the Euclidean distance between the five parameters diameter, circularity, cell wall thickness, cell wall distance from the PM and presence or absence of an OM for five replicate sets of measurements per sample (data from 5 micrographs and 3 - 5 tomograms per sample).

We assessed the robustness and specificity of the measurements, by comparing the similarity between replicates to the similarity between samples as given by the analysis of similarities (ANOSIM) R. Replicate measurements exhibited a high degree of similarity (R = 0.888, p < 10^−4^), indicating that our set of five discrete parameters achieves a meaningful separation between samples.

### The cellular architecture entails the ecology of carbon source utilization

To evaluate the sensitivity of the five parameters describing the cellular architecture, we evaluated whether they could detect changes in morphology in response to a change in carbon source. Because as environmental changes are reflected in protein expression profiles ^11^, we hypothesized that they might also be reflected in the cellular architecture.

We cultivated the human gut bacteria *Bacteroides thetaiotaomicron* DSM 2079 and *Ruminococcus bromii* L2-63, as well as the environmental isolate *Clostridium thermocellum* DSM 1313 either on a simple carbon source, i.e., glucose, fructose or cellobiose, or their preferred polysaccharide starch, pullulan or cellulose. The cellular architecture was quantified using five parameters as described above, followed by principal component analysis (PCA) across all measurements.

This analysis indicated that changes in carbon source correspond to overall structural variations for all the three strains that were analyzed pointing to a connection between ecology and cellular architecture (Fig. 4a). Interestingly, the structural response differs between the species. In the case of *B. thetaiotaomicron* DSM 2079, a change from glucose to starch/maltose was associated with a thicker cell wall (Fig. 4b, 6.8 ± 0.8 nm vs. 10.8 ± 1.1 nm). On the other hand, *R. bromii* L2-63 grown on low-complexity carbon source fructose exhibited a significantly lower diameter than either pullulan or starch (Fig. 4c, 0.673 ± 0.06 um, 0.898 ± 0.07 um and 0.889 ± 0.06 um, respectively).

**Figure 4:**
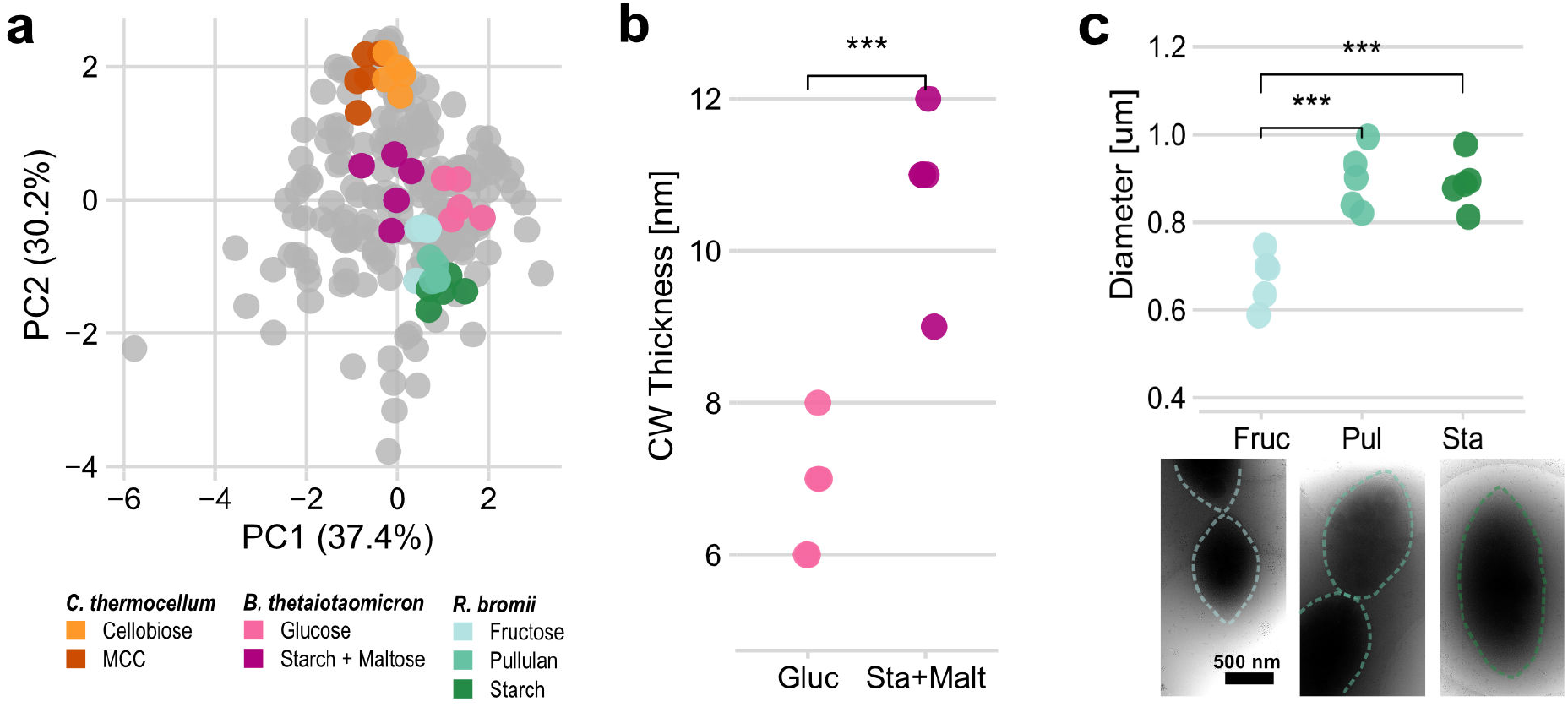
Cellular architecture reflects changing carbon sources. (a) Principal Component Analysis (PCA) of the five parameters diameter, circularity, presence of an outer membrane, cell wall thickness and cell wall distance from the plasma membrane across all 45 samples in the screening collection, of which three-dimensional data could be acquired. Highlighted are measurements for strains *Clostridium thermocellum* DSM 1313, *Bacteroides thetaiotaomicron* DSM 2079 and *Ruminococcus bromii* L2-63 grown on simple or complex carbon sources. (b) Further analysis shows that the cell wall of *Bacteroides thetaiotaomicron* DSM 2079 is significantly thicker when grown on starch/maltose (unpaired t-test, p < 0.001). (c) For *Ruminococcus bromii* L2-63, the cell diameter is higher when grown on complex carbon sources starch or pullulan compared to cells grown on fructose (unpaired t-tests, both p < 0.001). Example micrographs shown below.

The alterations in morphology of *C. thermocellum* DSM 1313 were non-significant for each individual structural parameters that were measured. The spread observed in the PCA might thus represent a combination of more subtle adaptations, such as the changes in the fiber degradation machinery we described in an earlier publication ^11^.

### Structural diversity arises in the phylogenetic branches

After evaluating the specificity and sensitivity of the similarity score based on five structural parameters, we asked whether the cellular architecture of a strain is linked to its phylogeny. Through ANOSIM, we assessed that replicates within the same strain or species show high degrees of structural similarity (R = 0.867, p < 10^−4^ and R = 0.854 respectively, both p < 10^−4^), some of which seems to be conserved within a genus (R = 0.711, p < 10^−4^). However, in-group similarity seems to be negligible at the family level (R = 0.242, p < 10^−4^) and above. To confirm this trend, we compared the Euclidean distance between each set of parameters to the lowest taxonomic rank they shared (Fig. 5, left). This analysis revealed that cells belonging to the same species usually have very similar cellular architectures, which increasingly differ for samples sharing only genus or family. Representative tomogram slices of three *Bacteroides* strains illustrate a conserved cellular architecture within a genus. On the contrary, three members of different genera in the family *Lachnospiraceae* exhibit distinct morphologies (Fig. 5, right). Interestingly, the link between shared taxonomic rank and structural similarity breaks above the family level; two strains belonging to the same order are as similar as those sharing only the domain.

**Figure 5:**
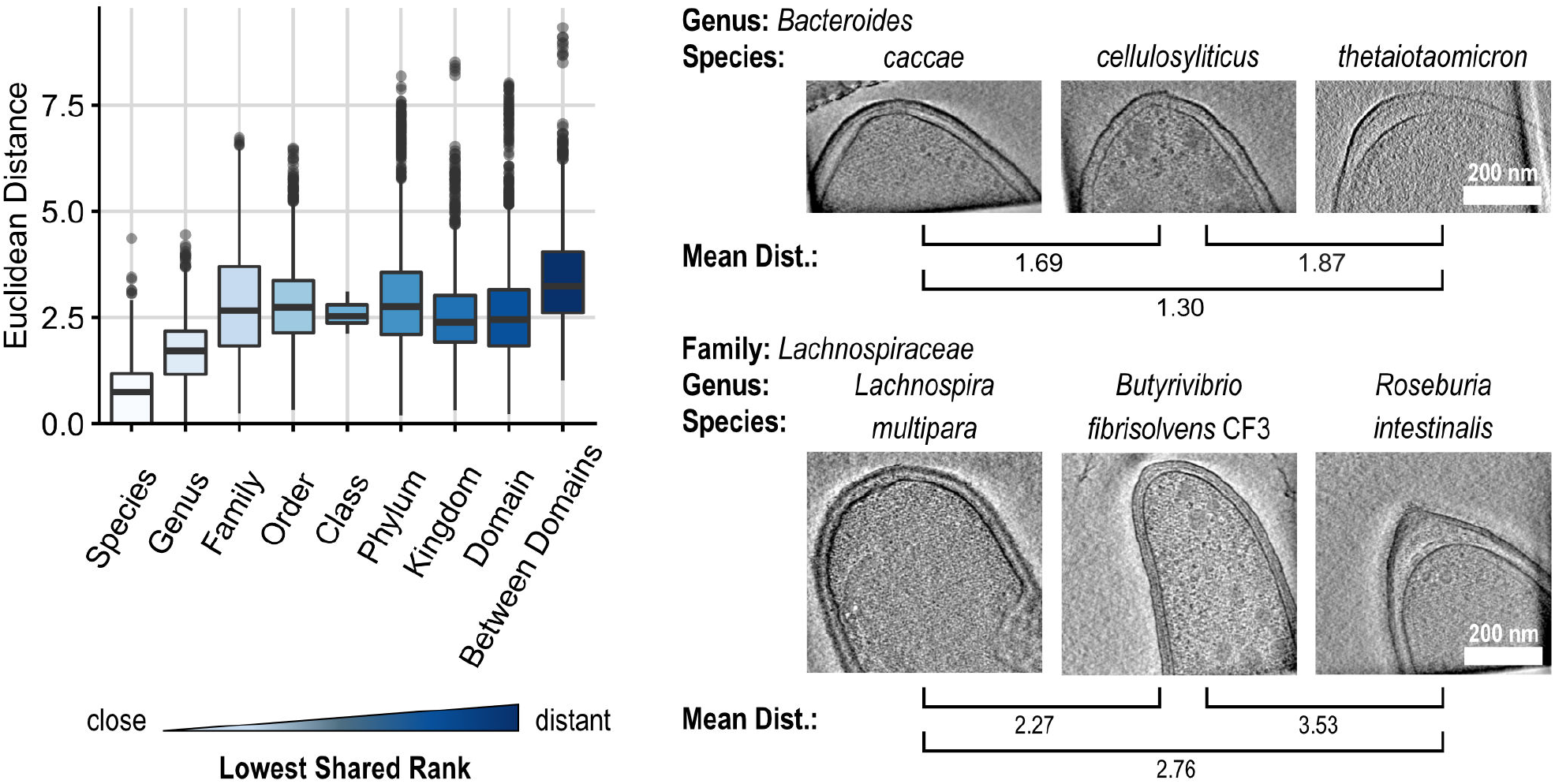
Visual Diversity emerges in the Phylogenetic Branches. The Euclidean distance between each pair of replicates as a function of their lowest shared taxonomic rank. Samples within a species show a conserved cellular architecture, as do those within a genus. Starting at the family level, the cellular architectures do not become more diverse within the same domain.

From this, we concluded that the distinct cellular architectures of microbiome members arises at the taxonomic levels below order, representing recent specialization rather than ancient branching events. Indeed, the structural similarity correlates significantly with the genomic distance for all pairs which share the species, genus or family (Pearson’s r = 0.68, p < 2.2 10^−16^), but not for those more distantly related (r = -0.01, p = 0.07).

As an example, tomogram slices from three representative strains in the genus *Bacteroides* are shown on the top right: *Bacteroides caccae* DSM 19024, *Bacteroides cellulosyliticus* DSM 14838 and *Bacteroides thetaiotaomicron* DSM 2079. All three present as rod-shaped diderm bacteria with thin cell walls. Correspondingly, the mean Euclidean distances between them are low. In contrast, three members of the *Lachnospiraceae* family show little resemblance and correspondingly higher Euclidean distances: *Lachnospira multipara* ATCC 19207, *Butyrivibrio fibrisolvens* CF3 and *Roseburia intestinalis* DSM 14610.

### In-family heterogeneity hints at diverse ecological roles

To better understand the broken link between taxonomy and cellular architecture, we closely inspected the taxonomy, genome and morphology of the most abundant family in the dataset, the *Lachnospiraceae*. Hierarchical clustering by genome contents and cellular architecture revealed that two strains belonging to the same genus have similar genetic compositions and similar cell architectures (Fig. 6a). However, the relationships between the genera are not conserved between genetic and morphological clustering.

**Figure 6:**
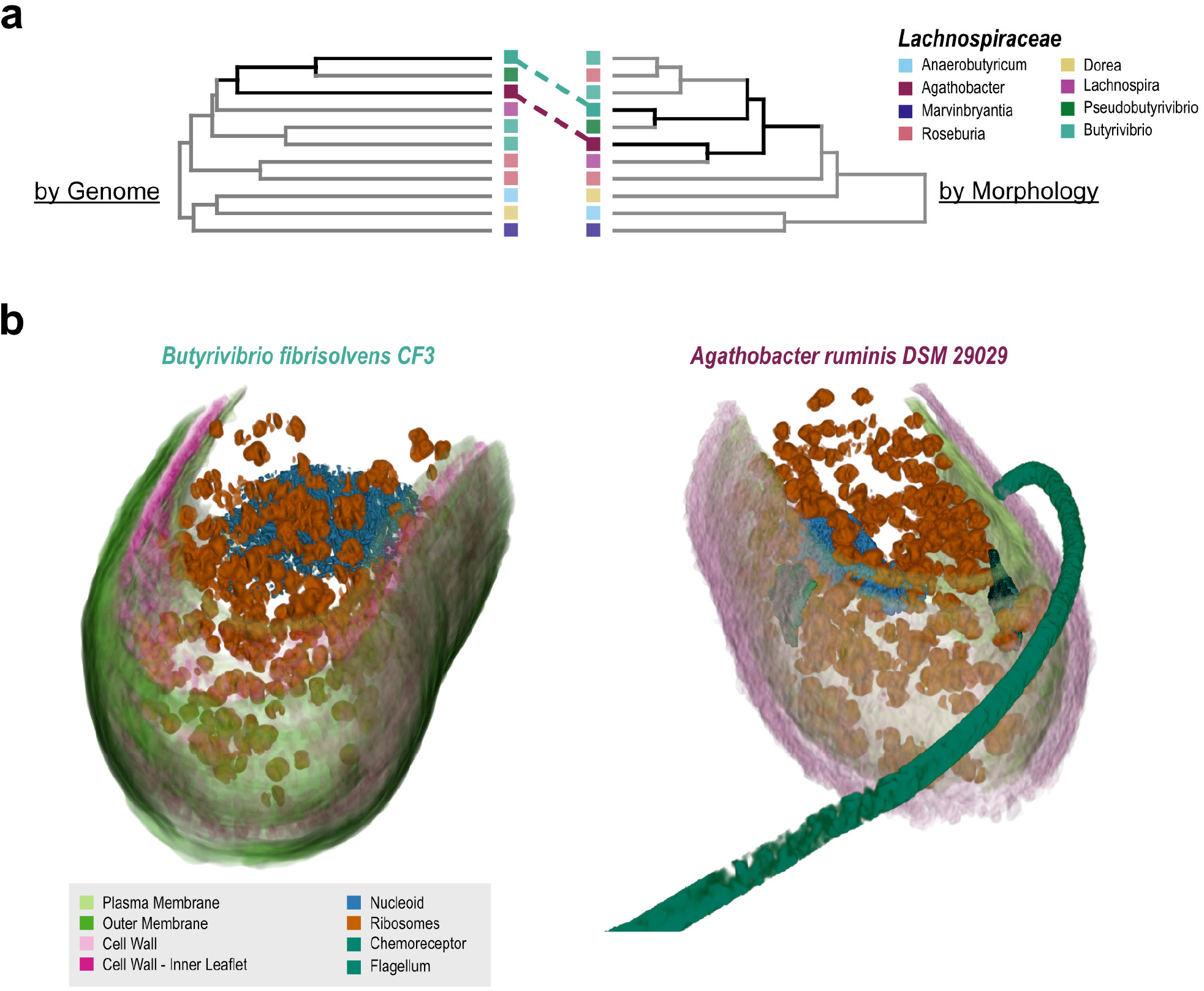
Hierarchical clustering and 3D rendering reveal divergence between genome and morphology. (a) All members of the *Lachnospiraceae* family were hierarchically clustered either by genome contents or by cellular architecture. Members of the same genus are clustered together in most cases, indicating a reliable link between taxonomy, genome and morphology at this level. The relationships between genera are shuffled: The example case of *Butyrivibrio fibrisolvens* CF3 and *Agathobacter ruminis* DSM 29029 is highlighted in both dendrograms. Corresponding samples are connected with dotted lines, relationships highlighted in black. (b) 3D renderings of *B. fibrisolvens* and *A. ruminis* cells. The divergent cell architectures are immediately visible: *B. fibrisolvens* is bound by a double-layered cell wall (pink) and an outer membrane (dark green), while *A. ruminis* is monoderm; it only carries one cell wall layer (pink) on top of the plasma membrane (light green). In both cells, the cytoplasm contains a high density of presumptive ribosomes (orange), as well as a nucleoid (blue). *A. ruminis* additionally contains two chemoreceptor arrays (teal), as well a flagellum in the extracellular space (teal).

We analyzed the implications of this divergence by visualizing the two *Lachnospiraceae* family members *Agathobacter ruminis* DSM 29029 and *Butyrivibrio fibrisolvens* CF3 in 3D (shift in dendrograms indicated by dotted lines in Fig. 6a). Based on the renderings, we could carefully inspect their diverging cellular architectures (Fig. 6b, supplementary videos 1 and 2).

While the diameter (*A. ruminis*: 0.377 ± 0.02 µm, *B. fibrisolvens*: 0.398 ± 0.01 µm) and circularity (0.407 ± 0.062 and 0.429 ± 0.108) were closely matched between these strains, *A. ruminis* is monoderm (Fig. 6, PM in light green and cell wall in pink), while *B. fibrisolvens* appears to be diderm (Fig. 6, OM in dark green) with a two-layered cell wall (similar to that of *Pseudobutyrivibrio sp*. in Fig. 3 a - c, its closest neighbor in both dendrograms), even though both are described as Gram-negative. Correspondingly, *A. ruminis* has a significantly thicker cell wall than *B. fibrisolvens* (18.8 ± 1.6 nm vs. 11.4 ± 0.8 nm, p < 0.001). In both cells, a high number of presumptive ribosomes (Fig. 6, orange) could be observed around a nucleoid (blue). In *A. ruminis*, a flagellum was observed connected to the cell, along with two chemoreceptor arrays (both teal) adjacent and opposite to the observed flagellar attachment site.

The unique features of both strains are well-known examples of structure-function relationships in microbiomes: flagella provide motility and access to distant nutrients ^8^ and capsules provide protection against phagocytosis ^9^. These divergent attributes in closely related strains could thus be the outcome of ecological constraints.

## Discussion

In this study, we used a combination of genomics, ecology and structural analysis to tackle the question of structure - function relationship in microbiome ecosystems. We studied a large collection of microbes that covers a substantial part of the diversity of the core rumen microbiome. Our work provides a general insight into variations of the cellular architecture of gut microorganisms. We show that a set of five cell parameters (diameter, circularity, presence of an OM, CW thickness and CW - PM distance) can be used to define the cellular architecture of a strain with a high specificity for 40 tested strains. While taxonomy and morphology are correlated below the family level, they diverge afterwards. We argue that structure is likely linked to ecological function and niche and thus selecting structurally diverse strains may be advantageous when reconstituting a representative ecosystem.

The application of cryo-EM and -ET allowed us to study a large number of bacterial and archaeal cells in a near-native state with nanometer resolutions to determine their cellular architecture. Cryo-EM provides detailed 2D description of individual cells, enriched by cryo-ET with 3D information on the organization of the cell wall and membrane, as well as the crowded cell interior. Our data complements a collection of bacterial and archaeal tomograms that were published by Jensen and coworkers (https://etdb.caltech.edu). We hope this publication provides a similarly useful collection by providing 3D information on an additional 40 strains to the public (an increase of 55%).

We first focused our attention to the macromolecular complexes visible in the tomograms. We frequently observed large and distinct structures such as ribosomes, storage granules, chemoreceptor arrays and flagellar motors. However, in tomograms in which we did not observe many of these complexes, we could not ascertain whether this represented biological ground truth rather than imaging constraints. A stark illustration of that is in Fig. 3. Even in the very thin *Pseudobutyrivibrio sp*. LB 2011, a dense cytoplasm completely obscures the electron-dense ribosomes. To compare cell morphologies across species, we thus decided to parametrize only the cell surface structure, which we could clearly image across all species. Likely, using more detailed descriptions of the molecular sociology of the cell could further elucidate the link between genotype and cellular architecture for Prokaryotes and Archaea, as well as their environmental role. Chemotaxis has been shown to shape microbial ecosystems ^25^ and flagella are both a virulence factor and trigger immune homeostasis in gut microbiomes ^26^.

Even though the five parameters (diameter, circularity, OM presence, cell wall thickness and distance from the PM) we used to quantify cellular architecture describe only the lowest common denominator of features visible in all tomograms, the structural similarity measure they provide was robust and sensitive as indicated by ANOSIM analysis. However, we propose that they should be regarded as reference values to detect changes in response to the cellular environment and not as a universal constant. Recent studies have shown that even fundamental features such as the cell wall can be remodeled or even lost completely under extreme circumstances ^27^. We indeed show that minor remodeling can also happen in response to a change in the environment of a microbe such as the type of carbon source under otherwise identical conditions for two prominent human gut microbiome members.

When comparing microbes’ structural similarity to the lowest common taxonomic rank, we observed a steady decrease in similarity for further relationships up to the family level, above which no link is discernible. Remarkably, this pattern was also observed for metabolic similarities by Han *et al*. ^28^. A preliminary analysis of the four strains included both in our study and their metabolomics dataset shows a high degree of correlation between the cellular architecture and metabolomic phenotype as indicated by a Mantel test, albeit not reaching statistical significance of 0.05 due to the limited number of possible permutations (r = 0.885, p = 0.08).

Overall, our findings suggest that the current taxonomic system does not sufficiently convey morphological information above the genus level. This underlines the importance of characterizing strains’ function individually and not taking taxonomies as shortcuts when comparing microbiome composition across individuals or ecosystems.

Applying structural analysis to a collection of microorganisms that represents much of the diversity within the core rumen microbiome opens an additional vantage point to study microbial relationships.

Microbial strains resembling those studied here are also present in other gut ecosystems, such as the human gut microbiome ^29,30^. Therefore, this dataset may serve as a basis to better understand structure-function relationships among microorganisms and enable future structural studies of reconstituted multi-species microbiomes. We observed a divergence between taxonomy and cellular architecture, which highlights challenges with phylogeny-based clustering of microorganisms. The preliminary link between metabolomics and morphology identified here in a small subset studied warrants further exploration. If it holds true that structural phenotype and metabolic function are correlated, it will prove a big step forward towards understanding the structure-function relationship of microbiomes. Ultimately, we imagine a classification system for microorganisms, which does not only take 16S-derived phylogenetic information into account, but could also encompass metabolomic information and high-resolution structural information from cryo-EM/ET. Such a classification system will help advance our understanding of the relationships within important microbial ecosystems.

## Methods

### Bacterial Culture

A collection of bacterial strains was assembled from commercially available culture collections (DSM and ATCC) or obtained from collaborations with USDA and the Rumen Microbial Genomics Network ^31^. The 69 bacterial strains were cultivated on their preferred media (Supplementary Table 1). Besides Brain Heart Infusion broth (Merck) and MRS (BD) that were purchased commercially, the various media were prepared as described previously, and included YCFA ^32^, M2 ^33^, BY ^34^ and GS2 ^35^. All media were supplemented with 1% (w/v) appropriate carbon sources as indicated in Supplementary Table 1. The bacterial cells were grown anaerobically at 37°C for 24h to 48h and the archaeal strains were incubated at 37°C until methane production was measurable. A volume 1ml of the cultures were harvested 5 min at 5000 *g*, washed briefly in TBS buffer, and resuspended in TBS prior to plunge-freezing. In parallel, 16S rRNA Sanger sequencing served to confirm the purity of the microbial cultures.

### Whole-Genome Sequencing

The genomic DNA of *Agathobacter ruminis* DSM 29029, *Bacteroides cellulosilyticus* DSM 14838, *Bacteroides thetaiotaomicron* DSM 2079, *Bacteroides vulgatus* ATCC 8482, *Bacteroides caccae* JCM 9498, *Bacteroides uniformis* ATCC 8492, *Butyrivibrio fibrisolvens* CF3, *Blautia schinkii* DSM 10518, *Butyrivibrio* sp. DSM 10294 and *Bifidobacterium thermophilum* DSM 20209 were extracted using the phenol-chloroform method ^36^. The purified DNA was sequenced using Illumina sequencing NovaSeq S1 and the obtained sequences were assembled using SPAdes ^37^. Genomes assemblies were deposited in GenBank (accession number SAMN30220820 to SAMN30220829).

### Cryo-EM Sample Preparation

Samples were prepared by plunge-freezing into liquid ethane. 5 µl of bacteria diluted to OD = 0.15 in PBS were deposited on a glow-discharged Quantifoil R2/1 holey carbon grids (Quantifoil Micro Tools GmbH, Germany) and 3 µl gold particles (10 nm BSA tracer, Aurion) concentrated 5-fold by centrifugation and resuspension in PBS were added to the samples that were then manually blotted for four seconds and vitrified by rapidly plunging into liquid ethane using a home-built plunging apparatus. The frozen samples were stored in liquid nitrogen until imaging.

### Cryo-EM Data Acquisition

Grids carrying frozen-hydrated samples were clipped into AutoGrids (TFS) and loaded into a Titan Krios microscope (FEI) equipped with a Gatan imaging filter and K2 direct electron detector. Data acquisition was controlled by SerialEM 3.8 ^38^. All micrographs were collected in gain-corrected counting mode with a slit width of 20 eV. Overview montages at a magnification of 4,800x (calibrated pixel size 3.4 nm) with defocus target -50 µm were used to measure diameter and shape of bacteria and guide further acquisition.

For suitably thin species, tilt series of dose-fractionated movies were acquired at 64,000x (calibrated pixel size 0.221 nm) in a bidirectional scheme starting at -30° spanning tilts ±60° in 3° increments with -4 - -6 µm underfocus. For *Ruminococcus bromii*, tilt series were acquired in a dose-symmetric scheme from 0° to ±60° at 105,000x magnification (pixel size 0.137 nm) ^39^.

For species judged too dense for tilt series acquisition, single projections were acquired at a magnification of 11,500x (pixel size 1.30 nm) as dose-fractionated movies at a target defocus of -8 µm with a total dose of ∼ 20 e^−^/A^2^.

Data for *Bacteroides thetaiotaomicron* DSM 2079 on glucose, *Bacteroides uniformis* ATCC 8492 and *Corynebacterium vitaeruminis* Ga6A13 were collected using a Titan Krios G3i system (TFS) equipped with a Gatan imaging filter and a K3 direct electron detector (Gatan). Imaging parameters were chosen to parallel those on the original system: Overview montages were acquired at 3,600x magnification (pixel size 2.4 nm), tilt series were collected at 42,000x magnification (pixel size 0.216 nm).

The projection dose was calibrated for each sample to yield a total dose of ∼140 e^−^/A^2^ for the tilt series (240 e^−^/A^2^ for *R. bromii*).

### Cryo-EM Data Processing

Cryo-EM data was processed using the *imod* software package ^40^. Montages were stitched using the tool *blendmont*, dose-fractionated movies were aligned using *alignframes* or the SerialEM SEMCCD plug-in. Tilt series were aligned, CTF-corrected by phase flipping, dose-weighted and reconstructed via weighted back-projection in *etomo*. For figure preparation, tomograms were deconvolved using tom_deconv.m ^41^.

### Imaging Data Analysis

Micrographs and Tomograms were opened in FIJI using the Bio-Formats Importer plugin ^42^. Distance measurements were performed using the line measurement tool. Shape quantification was performed as follows: Image thresholds were set so the cell wall or outer membrane was clearly contrasted. Then, the boundary of the microbe was traced using the polygon selection tool. A spline was fit through the points. Then, the shape was measured, and shape descriptors were extracted. Circularity, defined as 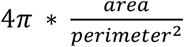 was used as a shape descriptor.

Measurement values are reported as mean and standard deviation of 5 - 10 individual measurements for diameter and circularity or 5 measurements in 3 - 5 tomograms for plasma membrane - cell wall distance and cell wall thickness.

### Statistical Analysis

Statistical analysis was performed in R (version 4.2) ^43^. Phylogenetic information was parsed from the NCBI taxonomy browser using the package taxize 0.9.99 ^44^. Distance matrices and ANOSIM were calculated using vegan 2.5-7 ^45^. For each bacterial strain, putative ORFs were predicted using Prokka ^46^ and then Proteinortho ^47^ was used to compare ortholog groups from all strains. Genomic distances were calculated using Jaccard indices. Hierarchical clustering was performed using hclust and exported via ape 5.6-1 ^48^ for visualization in itol ^49^.

Single-parameter comparisons for Fig. 4 and Fig. 6 were calculated using GraphPad Prism 9.3.1 (GraphPad Software, San Diego, USA). A Welch t-test was used with false discovery rate correction (Q = 1%) according to Benjamini, Krieger and Yekutieli.

To compare cellular architecture to metabolomic profile, the Metabolomics dataset published by the Sonneburg lab ^28^ was downloaded from Zenodo. The four strains included in the preliminary analysis are *Marvinbryantia formatexigens* DSM 14469, *Bacteroides cellulosilyticus* DSM 14838, *Bacteroides caccae* ATCC 43185 and *Eubacterium siraeum* DSM 15702.

### Data and Code Availability

The tomograms underlying the main text figures have been uploaded to the Electron Microscopy Data Bank (EMDB), accession codes EMD-15445 (*A. ruminis*), EMD-15447 (*B. fibrisolvens CF3*), EMD-15448 (*Pseudobutyrivibrio sp*.) and EMD-15449 (*Wolinella sp*.). Data and R code used for the statistical analysis are available on GitHub under https://github.com/bwmr/microbiome_morphology. Images of all samples in Supplementary Figure 2 are available on Zenodo under doi: 10.5281/zenodo.6874894.

Genomes assemblies were deposited in GenBank (accession number JANSWD000000000-JANSWM000000000).

## Acknowledgements

This work was funded by the DIP (2476/2 -1) to I.M. and O.M., ERC (866530) to I.M., ISF (1947/19) to I.M. and S.M. and the Swiss National Foundation (31003_207453) to O.M.. The authors thank Dr Meltem Tatli for providing the micrographs and tomograms of *C. thermocellum* on MCC and cellobiose and Daphne Perlman for assisting with graphic design. We acknowledge bacterial samples kindly provided by the USDA, the Hungate Collection (Dr Graeme T Attwood and Kerri Reilly), the Rumen Microbial Genomics Network, Claudia Moresi (ETH Zurich, *Bacteroides thetaiotaomicron* DSM 2079 on BHI-Hemin) and Prof. Magnus Øverlie Arntzen (NMBU Norway, *Fibrobacter succinogenes* S85 glycerol stocks). Cryo-electron microscopy imaging was performed with equipment maintained by the Center for Microscopy and Image Analysis, University of Zurich.

## Supplementary Figure Legends

**Supplementary Figure 1**

Phylogenetic tree of the rumen core microbiome and microbial collection examined in this study. The tree is based on the 16S rRNA gene of core microorganisms present in 90% of the individuals at the order level in a cohort of 1000 cows. Color coding is according to order level. Colored branches indicate the isolates examined in this study. Bootstraps with a confidence higher than 95% are displayed as pink circles. Samples with full 3D information colored black, if only micrographs were available in gray. The tree was created using iTOL.

**Supplementary Figure 2**

Exemplary medium-magnification micrographs and 1 nm thick tomogram slices for all samples in the dataset. The outline of the microbe is traced in the micrograph as used for circularity measurements. List is sorted alphabetically. TIF versions of all images are available on Zenodo, doi: 10.5281/zenodo.6874894

**Supplementary Table 1**

Average measurements and standard deviations for all samples in the dataset.

**Supplementary Movies for Figure 6**

- **<u>Supplementary Movie 1:</u>** <u>Tomogram and rendering of</u> *<u>Butyrivibrio fibrisolvens</u>* <u>CF3.</u>
- **<u>Supplementary Movie 2</u>**<u>: Tomogram and rendering of</u> *<u>Agathobacter ruminis</u>* <u>DSM 29029.</u>

