## Supplementary File for "A link between genotype and cellular architecture in microbiome members as revealed by cryo-EM"

Contents**:**

- **Supplementary Table 1**: Average measurements and standard deviations for all samples in the dataset.

- **Supplementary Figure 1:** Phylogenetic tree of the microbial collection examined in this study.

- **Supplementary Figure 2:** Representative micrographs and tomogram slices of all samples in this study.

| | Sample Information | | | | cryo-EM Quantifications | | | | | | | | | | | | |
|---|---|---|---|---|---|---|---|---|---|---|---|---|---|---|---|---|---|
| | | | | | From 2D Micrographs | | | | | | From 3D Tomography Data | | | | | | |
| Species / Strain | Identifier | Growth Medium | Isolation Host | Gram Staining | Diameter Mean [µm] | SD Diameter [µm] | Circularity | SD Circularity | Shape | n (cells) | Cell Surface [1] | Distance PM - CW [nm] | SD PM - CW | Thickness CW [nm] | SD CW | n (tomograms) | Sequenced for this study |
| **Acetivibrio_thermocellus_LQ8_Clostridiu** | DSM_1313 | GS2 + Cellobiose | Environmental | positive | 0.497 | 0.043 | 0.517 | 0.074 | Rod | 10 | Monoderm | 17.5 | 3.5 | 8.8 | 0.7 | 5 | |
| **Acetivibrio_thermocellus_LQ8_Clostridiu** | DSM_1313 | GS2 + cellulose | Environmental | positive | 0.534 | 0.046 | 0.375 | 0.077 | Rod | 10 | Monoderm | 18.4 | 3.1 | 11.8 | 2.3 | 5 | |
| **Actinomyces_ruminicola_B71** | DSM_27982 | M2+starch+cellobiose+glucose | Cow | positive | 0.630 | 0.055 | 0.673 | 0.094 | Rod | 10 | *Monoderm* | | | | | | |
| **Agathobacter_ruminis** | DSM_29029 | YCFA+ Glucose | Sheep | negative | 0.377 | 0.020 | 0.407 | 0.062 | Vibroid | 10 | Monoderm | 21.2 | 2.4 | 18.8 | 1.6 | 5 | yes |
| **Akkermansia_muciniphila_Muc** | DSM_22959 | YCFA+ Glucose + Mucin | Human | negative | 0.816 | 0.065 | 0.914 | 0.051 | Rod | 10 | Diderm | 16.4 | 1.0 | 13.6 | 1.6 | 5 | |
| **Anaerobutyricum_hallii** | DSM_3353 | YCFA+ Glucose | Human | positive | 1.122 | 0.094 | 0.621 | 0.065 | Rod | 10 | Monoderm | 27.8 | 4.4 | 27.2 | 3.1 | 4 | |
| **Anaerovibrio_lipolyticus_5S** | DSM_3074 | YCFA+ Glucose | Sheep | negative | 0.593 | 0.051 | 0.519 | 0.058 | Vibroid | 10 | Diderm | 20.4 | 3.6 | 10.4 | 2.4 | 5 | |
| **Bacteroides_caccae** | DSM_19024 | YCFA+ Glucose | Human | negative | 0.782 | 0.053 | 0.816 | 0.032 | Rod | 10 | Diderm | 25.6 | 1.9 | 12.4 | 1.0 | 5 | yes |
| **Bacteroides_cellulosyliticus_CRE21** | DSM_14838 | YCFA+ Glucose | Human | negative | 0.585 | 0.038 | 0.735 | 0.222 | Rod | 5 | Diderm | 20.8 | 2.7 | 8.8 | 1.0 | 3 | yes |
| **Bacteroides_thetaiotaomicron** | DSM_2079 | YCFA+ Glucose | Human | negative | 0.876 | 0.042 | 0.720 | 0.079 | Rod | 10 | Diderm | 19.2 | 1.9 | 6.8 | 0.7 | 3 | yes |
| **Bacteroides_thetaiotaomicron** | DSM_2079 | YCFA+ Starch / Maltose | Human | negative | 0.802 | 0.063 | 0.466 | 0.088 | Rod | 10 | Diderm | 27.2 | 3.6 | 10.8 | 1.0 | 4 | yes |
| **Bacteroides_thetaiotaomicron** | DSM_2079 | BHI + Cysteine + Hemin | Human | negative | 0.849 | 0.057 | 0.670 | 0.071 | Rod | 10 | Diderm | 23.8 | 2.9 | 8.6 | 1.0 | 3 | yes |
| **Bacteroides_uniformis** | ATCC_8492 | YCFA+ Glucose | Human | negative | 0.872 | 0.059 | 0.745 | 0.181 | Rod | 10 | *Diderm* | | | | | | yes |
| **Bacteroides_vulgatus** | ATCC_8482 | YCFA+ Glucose | Cow | negative | 0.956 | 0.060 | 0.736 | 0.120 | Rod | 10 | *Diderm* | | | | | | yes |
| **Basfia_succiniciproducens** | DSM_22022 | YCFA+ Glucose | Cow | negative | 0.942 | 0.111 | 0.816 | 0.072 | Rod | 10 | Diderm | 23.6 | 3.4 | 11.8 | 2.1 | 5 | |
| **Bifidobacterium_adolescentis** | DSM_20087 | YCFA+ Glucose | Cow | positive | 0.728 | 0.088 | 0.767 | 0.115 | Rod | 10 | Monoderm | 24.0 | 2.6 | 21.4 | 1.7 | 5 | |
| **Bifidobacterium_animalis_UR1** | DSM_10140 | YCFA+ Glucose | Yoghurt | positive | 0.760 | 0.035 | 0.583 | 0.083 | Rod | 10 | *Monoderm* | | | | | | |
| **Bifidobacterium_choerinum** | AGR2158 | YCFA+ Glucose | Cow | positive | 0.892 | 0.083 | 0.740 | 0.093 | Rod | 10 | *Monoderm* | | | | | | |
| **Bifidobacterium_ruminantium** | DSM_6489 | YCFA+ Glucose | Cow | positive | 0.699 | 0.039 | 0.726 | 0.088 | Rod | 10 | Monoderm | 23.0 | 1.0 | 21.5 | 1.5 | 5 | |
| **Bifidobacterium_thermophilum_C3-12** | DSM_20209 | YCFA+ Glucose | Ruminator, unspe | NA | 0.717 | 0.229 | 0.785 | 0.075 | Rod | 10 | *Monoderm* | | | | | | yes |
| **Blautia_schinkii** | DSM_10518 | YCFA+ Glucose | Lamb | positive | 0.907 | 0.037 | 0.748 | 0.042 | Rod | 10 | *Diderm* | | | | | | yes |
| **Butyrivibrio_fibrisolvens_CF3** | NA | YCFA+ Glucose | Cow | negative | 0.429 | 0.010 | 0.429 | 0.108 | Vibroid | 10 | Diderm | 17.0 | 2.6 | 11.4 | 0.8 | 3 | yes |
| **Butyrivibrio_fibrisolvens_D1** | DSM_3071 | YCFA+ Glucose | Cow | positive | 0.523 | 0.026 | 0.578 | 0.043 | Rod | 10 | Diderm | 28.2 | 1.0 | 6.8 | 1.7 | 5 | |
| **Butyrivibrio_sp** | DSM_10294 | YCFA+ Glucose | Cow | NA | 0.525 | 0.021 | 0.555 | 0.092 | Rod | 10 | Diderm | 22.0 | 0.6 | 5.0 | 1.1 | 3 | yes |
| **Clostridium_aminophilum_F** | DSM_10710 | YCFA+ Glucose | Cow | positive | 0.818 | 0.058 | 0.862 | 0.033 | Almond | 10 | Diderm | 19.4 | 4.0 | 7.0 | 1.4 | 5 | |
| **Clostridium_butyricum_5001_beijerinckii** | NA | YCFA+ Glucose | Cow | positive | 1.148 | 0.072 | 0.552 | 0.123 | Rod | 10 | *Diderm* | | | | | | |
| **Clostridium_cadaveris_G65** | AGR2141 | YCFA+ Glucose | Cow | positive | 0.728 | 0.038 | 0.627 | 0.073 | Rod | 10 | Monoderm | 29.0 | 1.9 | 24.8 | 1.0 | 5 | |
| **Clostridium_paraputrificum_G594** | AGR2156 | M2+starch+cellobiose+glucose | Cow | positive | 0.589 | 0.038 | 0.347 | 0.046 | Rod | 10 | Monoderm | 21.4 | 4.6 | 24.0 | 3.0 | 5 | |
| **Corynebacterium_vitaeruminis_Ga6A13** | NA | YCFA+ Glucose | Cow | positive | 0.995 | 0.054 | 0.778 | 0.061 | Rod | 10 | *Diderm* | | | | | | |
| **Dorea_sp** | AGR2135 | YCFA+ Glucose | Cow | positive | 0.670 | 0.060 | 0.667 | 0.084 | Almond | 10 | Monoderm | 34.2 | 3.2 | 33.6 | 2.9 | 4 | |
| **Eubacterium_pyruvativorans_KHPC4** | NA | M2+starch+cellobiose+glucose | Cow | positive | 0.672 | 0.052 | 0.727 | 0.095 | Rod | 10 | Monoderm | 22.8 | 1.6 | 20.6 | 3.4 | 5 | |
| **Eubacterium_siraeum** | DSM_15702 | YCFA+ Glucose | Human | positive | 0.645 | 0.079 | 0.783 | 0.121 | Rod | 10 | Monoderm | 19.2 | 1.9 | 34.8 | 3.2 | 5 | |
| **Faecalibacterium_prausnitzii_A2-165** | DSM_17677 | YCFA+ Glucose | Human | positive | 0.852 | 0.119 | 0.552 | 0.101 | Rod | 10 | Diderm | 18.6 | 1.0 | 10.6 | 1.2 | 4 | |
| **Fibrobacter_succinogenes_S85** | ATCC_19169 | YCFA - 0.5% Cellobiose | Cow | negative | 1.308 | 0.185 | 0.987 | 0.007 | Coccoid | 10 | *Diderm* | | | | | | |
| **Fusobacterium_necrophorum** | Hun48 | YCFA+ Glucose | Cow | negative | 0.819 | 0.099 | 0.540 | 0.106 | Rod | 10 | *Diderm* | | | | | | |
| **Kandleria_vitulina** | DSM_20405 | MRS (Glucose) | Cow | positive | 0.651 | 0.046 | 0.744 | 0.106 | Rod | 10 | Monoderm | 28.0 | 3.0 | 30.8 | 2.1 | 3 | |
| **Lachnobacterium_bovis** | DSM_14045 | YCFA+ Glucose | Cow | positive | 0.772 | 0.048 | 0.615 | 0.061 | Rod | 10 | *Monoderm* | | | | | | |
| **Lachnospira_multipara_G6** | ATCC_19207 | YCFA+ Glucose | Cow | positive | 0.460 | 0.019 | 0.479 | 0.090 | Rod | 10 | Monoderm | 29.6 | 3.5 | 22.8 | 3.7 | 5 | |
| **Ligilactobacillus_ruminis_Lactobacillus_** | DSM_20403 | MRS (Glucose) | Cow | positive | 0.623 | 0.035 | 0.497 | 0.143 | Rod | 10 | Monoderm | 38.6 | 9.5 | 53.6 | 6.7 | 5 | |
| **Limosilactobacillus_mucosae_Lactobaci** | AGR63 | MRS (Glucose) | Cow | positive | 0.726 | 0.011 | 0.631 | 0.119 | Rod | 10 | *Monoderm* | | | | | | |
| **Limosilactobacillus_reuteri_Lactobacillus** | DSM_17938 | MRS (Glucose) | Human | positive | 0.816 | 0.099 | 0.798 | 0.130 | Rod | 10 | *Monoderm* | | | | | | |
| **Marvinbryantia_formatexigens** | DSM_14469 | YCFA+ Glucose | Human | positive | 1.141 | 0.060 | 0.841 | 0.070 | Almond | 10 | Monoderm | 41.4 | 8.1 | 13.2 | 1.0 | 4 | |
| **Megamonas_sp_Calf98_2** | NA | YCFA+ Glucose | Cow | NA | 1.509 | 0.194 | 0.458 | 0.084 | Rod | 10 | *Diderm* | | | | | | |
| **Megasphaera_elsdenii_LC1** | DSM_20460 | YCFA+ Glucose | Sheep | negative | 1.805 | 0.083 | 0.983 | 0.015 | Coccoid | 10 | *Diderm* | | | | | | |
| **Methanobrevibacter_ruminantium_M1** | DSM_1093 | BY | Cow | NA | 0.613 | 0.045 | 0.629 | 0.120 | Vibroid | 10 | *Archeaum* | | | | | | |
| **Methanocorpusculum_labreanum_Z** | DSM_4855 | BY | Environmental | NA | 1.037 | 0.138 | 0.950 | 0.040 | Coccoid | 10 | Archeaum | 7.2 | 0.7 | 4.0 | 0.0 | 5 | |
| **Methanomicrobium_mobile_BP** | DSM_1539 | BY | Cow | NA | 0.737 | 0.042 | 0.837 | 0.070 | Almond | 10 | Archeaum | 8.2 | 1.0 | 4.6 | 0.5 | 5 | |
| **Micrococcales_bacterium_KH10** | NA | YCFA+ glucose | Cow | positive | 0.705 | 0.105 | 0.719 | 0.117 | Rod | 10 | *Diderm* | | | | | | |
| **Mitsuokella_jalaludinii_M9** | DSM_13811 | YCFA+ glucose | Cow | negative | 0.864 | 0.027 | 0.596 | 0.081 | Rod | 10 | *Diderm* | | | | | | |
| **Olsenella_umbonata_strain_WCP15** | NA | YCFA+ glucose | Cow | positive | 0.653 | 0.031 | 0.858 | 0.080 | Almond | 10 | *Monoderm* | | | | | | |
| **Pediococcus_acidilactici_N775** | AGR20 | YCFA+ glucose | Sheep | positive | 0.917 | 0.064 | 0.919 | 0.026 | Coccoid | 8 | *Monoderm* | | | | | | |
| **Peptostreptococcus_anaerobius_strain_(** | NA | YCFA+ glucose | Cow | positive | 0.921 | 0.058 | 0.951 | 0.048 | Coccoid | 10 | *Diderm* | | | | | | |
| **Prevotella_bryantii_B14** | DSM_11371 | YCFA+ glucose | Cow | negative | 0.964 | 0.043 | 0.600 | 0.077 | Rod | 10 | Diderm | 20.4 | 1.5 | 12.6 | 2.1 | 5 | |
| **Prevotella_ruminicola** | ATCC_19189 | YCFA+ glucose | Cow | negative | 0.693 | 0.044 | 0.868 | 0.044 | Almond | 10 | Diderm | 18.6 | 0.8 | 5.2 | 0.7 | 5 | |
| **Propionibacteriaceae_bacterium_P6A17** | NA | YCFA+ glucose | Cow | positive | 0.691 | 0.026 | 0.858 | 0.044 | Rod | 10 | Diderm | 10.4 | 0.5 | 7.4 | 0.5 | 5 | |
| **Proteiniclasticum_ruminis** | DSM_24773 | YCFA+ glucose | Yak | negative | 0.711 | 0.041 | 0.500 | 0.086 | Rod | 10 | Monoderm | 24.8 | 4.0 | 37.8 | 7.9 | 3 | |
| **Pseudobutyrivibrio_sp_LB2011** | NA | M2+starch+cellobiose+glucose | Cow | positive | 0.416 | 0.024 | 0.455 | 0.061 | Vibroid | 10 | Diderm | 14.6 | 0.8 | 13.4 | 0.8 | 5 | |
| **Roseburia_intestinalis_L1-82** | DSM_14610 | YCFA+ glucose | Human | positive | 0.663 | 0.054 | 0.367 | 0.053 | Rod | 10 | Monoderm | 40.0 | 10.8 | 6.0 | 1.1 | 5 | |
| **Roseburia_inulinivorans** | DSM_16841 | YCFA+ glucose | Human | positive | 0.613 | 0.039 | 0.613 | 0.065 | Rod | 10 | Diderm | 27.0 | 4.0 | 6.6 | 1.4 | 5 | |
| **Ruminococcus_albus_7** | DSM_20455 | M2+ cellobiose | Cow | positive | 1.206 | 0.116 | 0.955 | 0.029 | Coccoid | 10 | *Monoderm* | | | | | | |

| | Sample Information | | | | cryo-EM Quantifications | | | | | | | | | | | | |
|---|---|---|---|---|---|---|---|---|---|---|---|---|---|---|---|---|---|
| | | | | | From 2D Micrographs | | | | | | From 3D Tomography Data | | | | | | |
| Species / Strain | Identifier | Growth Medium | Isolation Host | Gram Staining | Diameter Mean [µm] | SD Diameter [µm] | Circularity | SD Circularity | Shape | n (cells) | Cell Surface [1] | Distance PM - CW [nm] | SD PM - CW | Thickness CW [nm] | SD CW | n (tomograms) | Sequenced for this study |
| **Ruminococcus_bromii_L2-63** | NA | M2+ starch | Human | positive | 0.863 | 0.036 | 0.887 | 0.041 | Almond | 10 | Monoderm | 25.4 | 2.6 | 13.4 | 2.2 | 5 | |
| **Ruminococcus_bromii_L2-63** | NA | M2+ fructose | Human | positive | 0.664 | 0.042 | 0.892 | 0.050 | Almond | 10 | Monoderm | 26.8 | 2.4 | 11.2 | 1.5 | 5 | |
| **Ruminococcus_bromii_L2-63** | NA | M2+ pullulan | Human | positive | 0.841 | 0.077 | 0.829 | 0.072 | Almond | 10 | Monoderm | 25.6 | 1.7 | 11.6 | 1.2 | 5 | |
| **Ruminococcus_champanellensis_18P13** | JCM_17042 | M2+ cellobiose | Human | positive | 0.849 | 0.029 | 0.869 | 0.065 | Almond | 10 | *Monoderm* | | | | | | |
| **Ruminococcus_champanellensis_18P13** | JCM_17042 | M2+ cellulose | Human | positive | 0.900 | 0.034 | 0.875 | 0.055 | Almond | 9 | *Monoderm* | | | | | | |
| **Ruminococcus_gnavus** | AGR2154_Hun2: | M2+starch+cellobiose+glucose | Cow | positive | 0.930 | 0.044 | 0.780 | 0.066 | Almond | 10 | *Monoderm* | | | | | | |
| **Selenomonas_bovis** | DSM_23594 | YCFA+ glucose | Yak | negative | 0.759 | 0.031 | 0.524 | 0.091 | Rod | 10 | Diderm | 18.0 | 0.9 | 9.2 | 1.9 | 3 | |
| **Selenomonas_ruminantium_AB3002_P15** | NA | YCFA+ glucose | Cow | negative | 0.975 | 0.070 | 0.586 | 0.068 | Rod | 10 | Diderm | 20.2 | 0.7 | 12.8 | 2.0 | 4 | |
| **Sharpea_azabuensis** | DSM_20406 | MRS (Glucose) | Cow | positive | 0.511 | 0.032 | 0.435 | 0.039 | Rod | 10 | Monoderm | 30.8 | 1.3 | 30.0 | 2.8 | 5 | |
| **Streptococcus_bovis** | ATCC_33317 | BHI | Cow | positive | 0.962 | 0.055 | 0.909 | 0.046 | Almond | 10 | *Monoderm* | | | | | | |
| **Streptococcus_henryi_A_4** | NA | YCFA+ glucose | Cow | positive | 0.829 | 0.067 | 0.964 | 0.012 | Coccoid | 10 | *Monoderm* | | | | | | |
| **Succinimonas_amylolytica** | DSM_2873 | YCFA+ glucose | Cow | negative | 1.374 | 0.126 | 0.994 | 0.003 | Coccoid | 10 | *Diderm* | | | | | | |
| **Succinivibrio_dextrinosolvens** | DSM_3072 | YCFA+ glucose | Cow | negative | 0.572 | 0.041 | 0.660 | 0.093 | Vibroid | 10 | Diderm | 16.4 | 2.6 | 9.0 | 1.4 | 5 | |
| **Terrisporobacter_glycolicus_strain_KPPI** | NA | YCFA+ glucose | Cow | positive | 0.861 | 0.040 | 0.765 | 0.100 | Rod | 10 | *Monoderm* | | | | | | |
| **Wolinella_sp_Vibrio_succinogenes** | ATCC_33567 | YCFA+ fumarate/formate/aspart | Cow | negative | 0.604 | 0.041 | 0.466 | 0.053 | Rod | 10 | Diderm | 21.0 | 6.0 | 12.2 | 3.4 | 5 | |

**Annotations**

[1] for samples without tomography, the mono- / diderm classification is lower confidence and thus marked in italics and grey.

### Supplementary Figure 1

Phylogenetic tree of the rumen core microbiome and microbial collection examined in this study. The tree is based on the 16S rRNA gene of core microorganisms present in 90% of the individuals at the order level in a cohort of 1000 cows. Color coding is according to order level. Colored branches indicate the isolates examined in this study. Bootstraps with a confidence higher than 95% are displayed as pink circles. Samples with full 3D information colored black, if only micrographs were available in grey. The tree was created using iTOL.

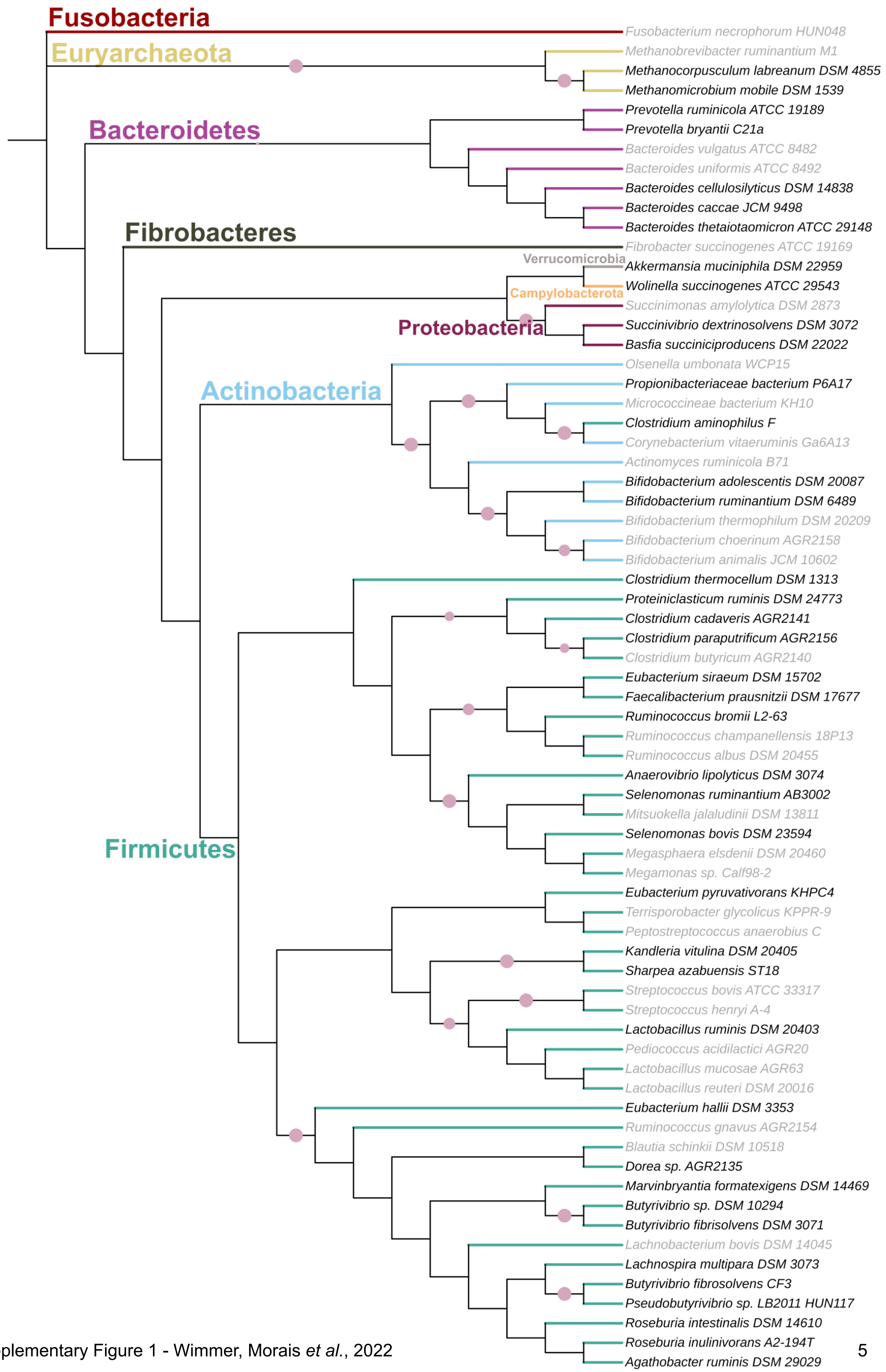


Supplementary Figure 1 - Wimmer, Morais *et al.*, 2022

### Supplementary Figure 2

Representative medium-magnification micrographs and 1 nm thick tomogram slices for all samples in the dataset. The outline of the microbe is traced in the micrograph as used for circularity measurements. List is sorted alphabetically. TIF versions of all images are available on Zenodo, doi: 10.5281/zenodo.6874894

### *Acetivibrio thermocellus / Clostridium thermocellum* DSM 1313

On Cellobiose

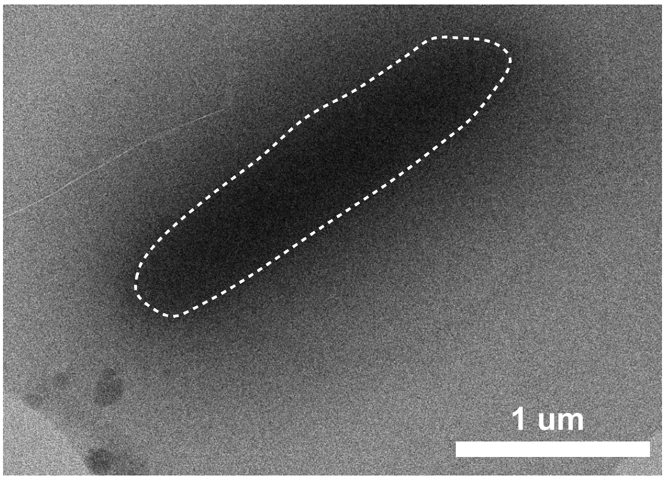


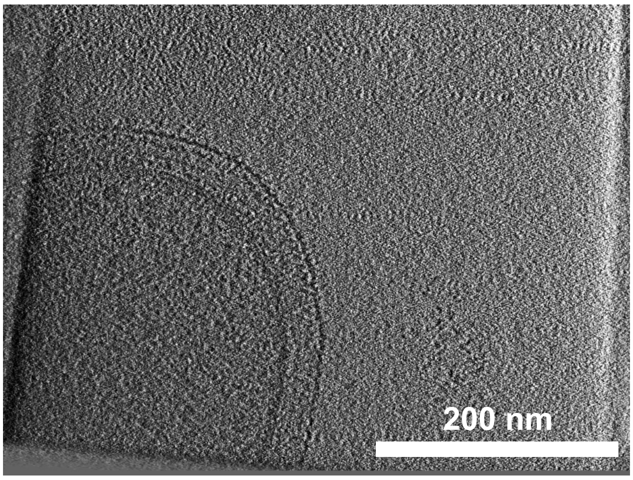


On MCC

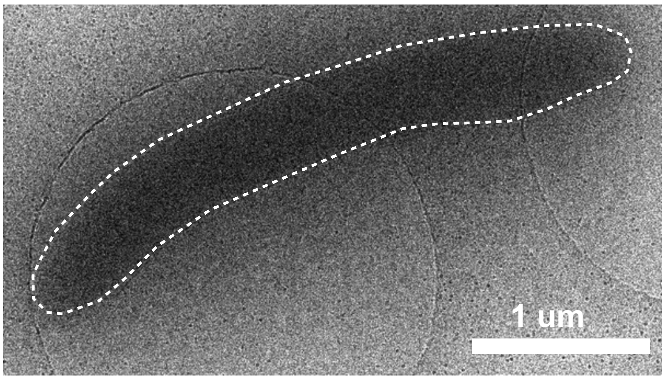


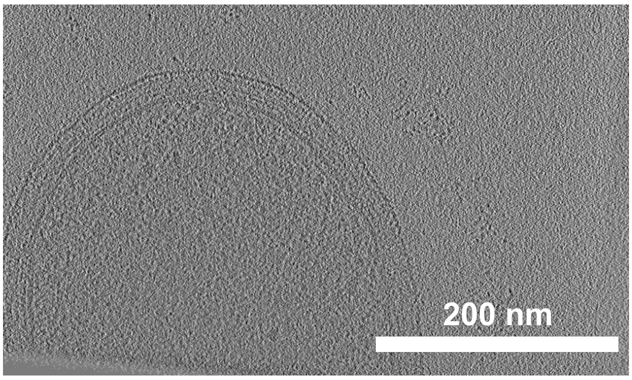


### *Actinomyces ruminicola* DSM 27982

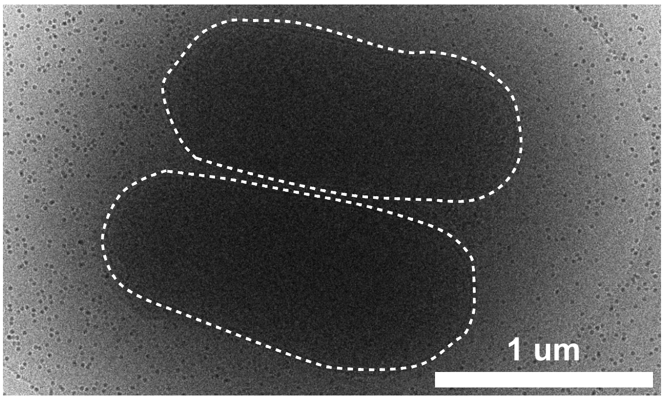


### *Agathobacter ruminis* DSM 29029

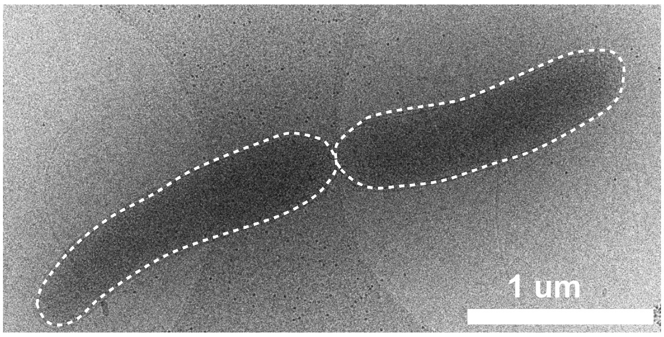


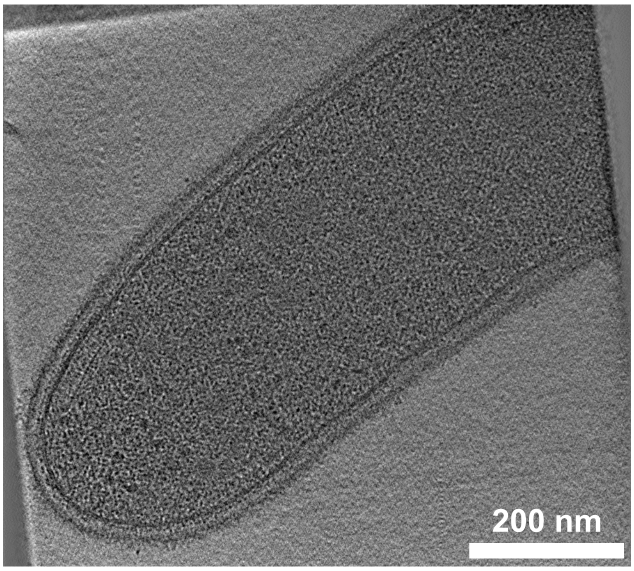

*Akkermansia muciniphila* DSM 22959

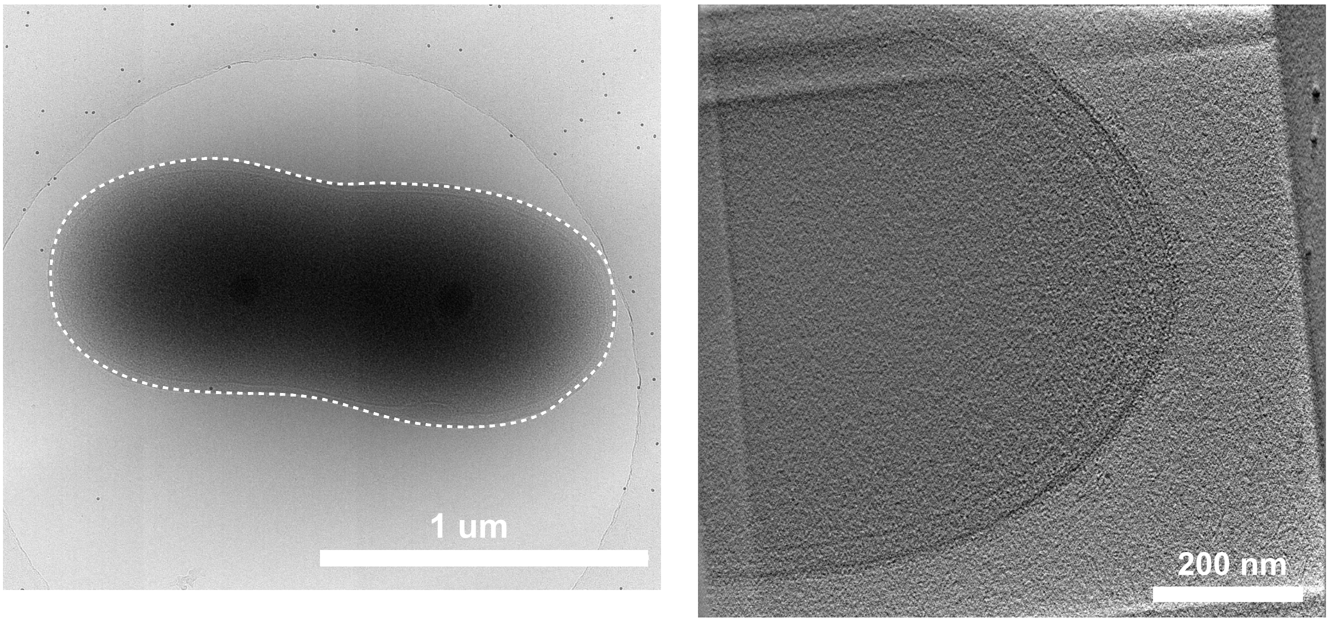


*Anaerobutyricum hallii* DSM 3353

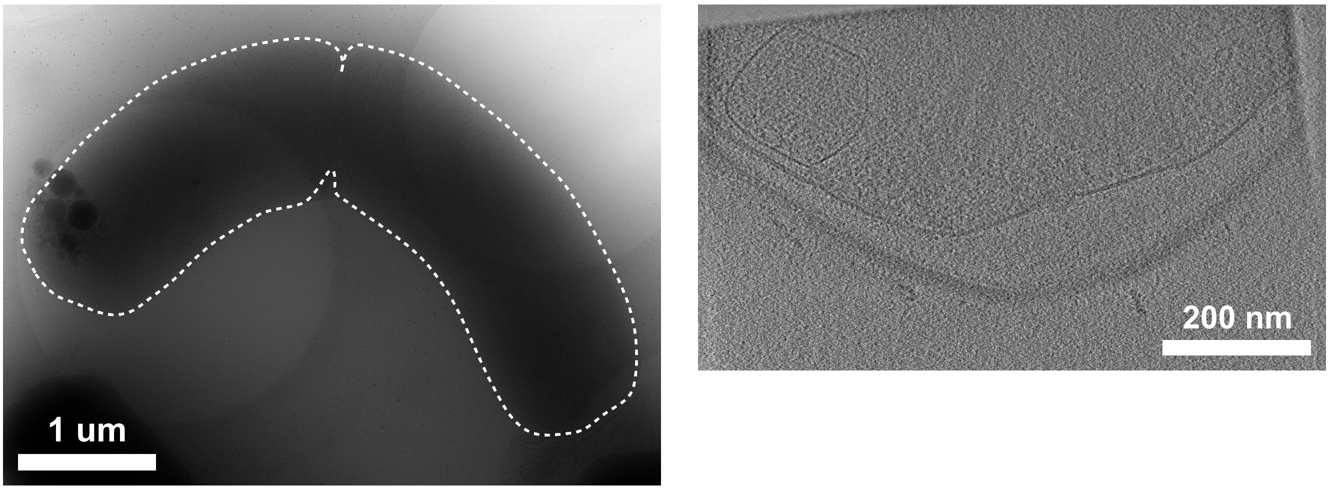


*Anaerovibrio lipolyticus* DSM 3074

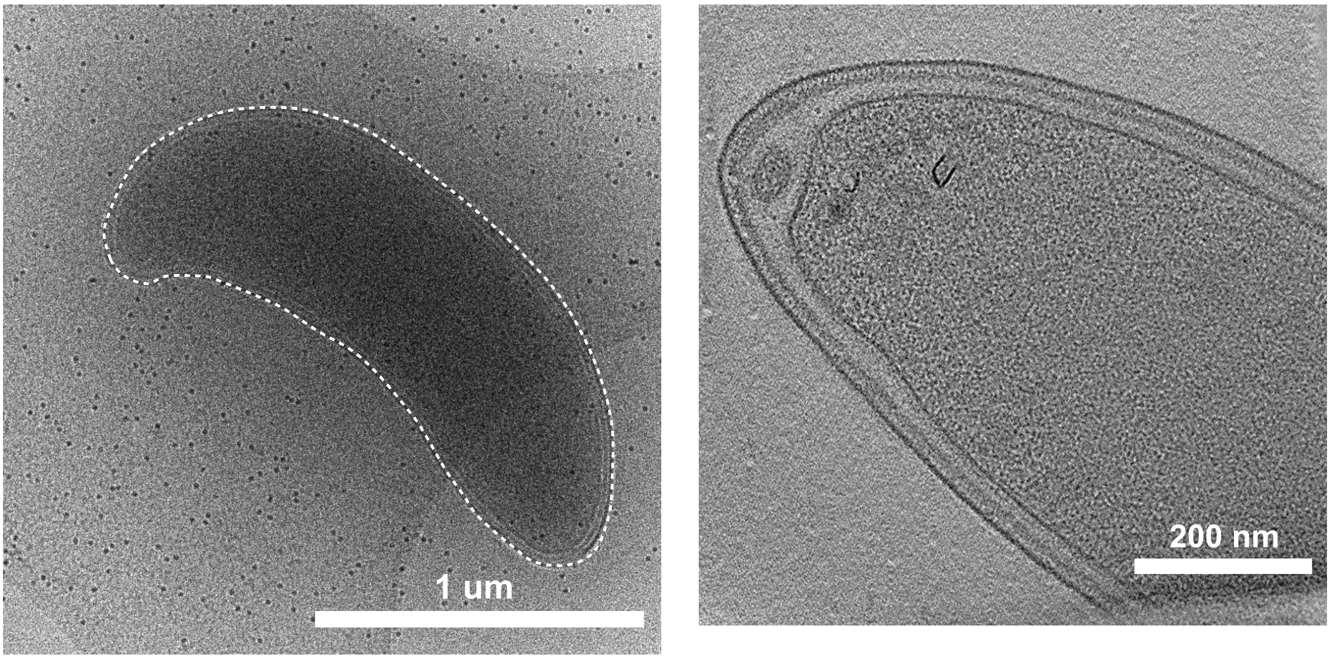

<u>*Bacteroides caccae* DSM 19024</u>

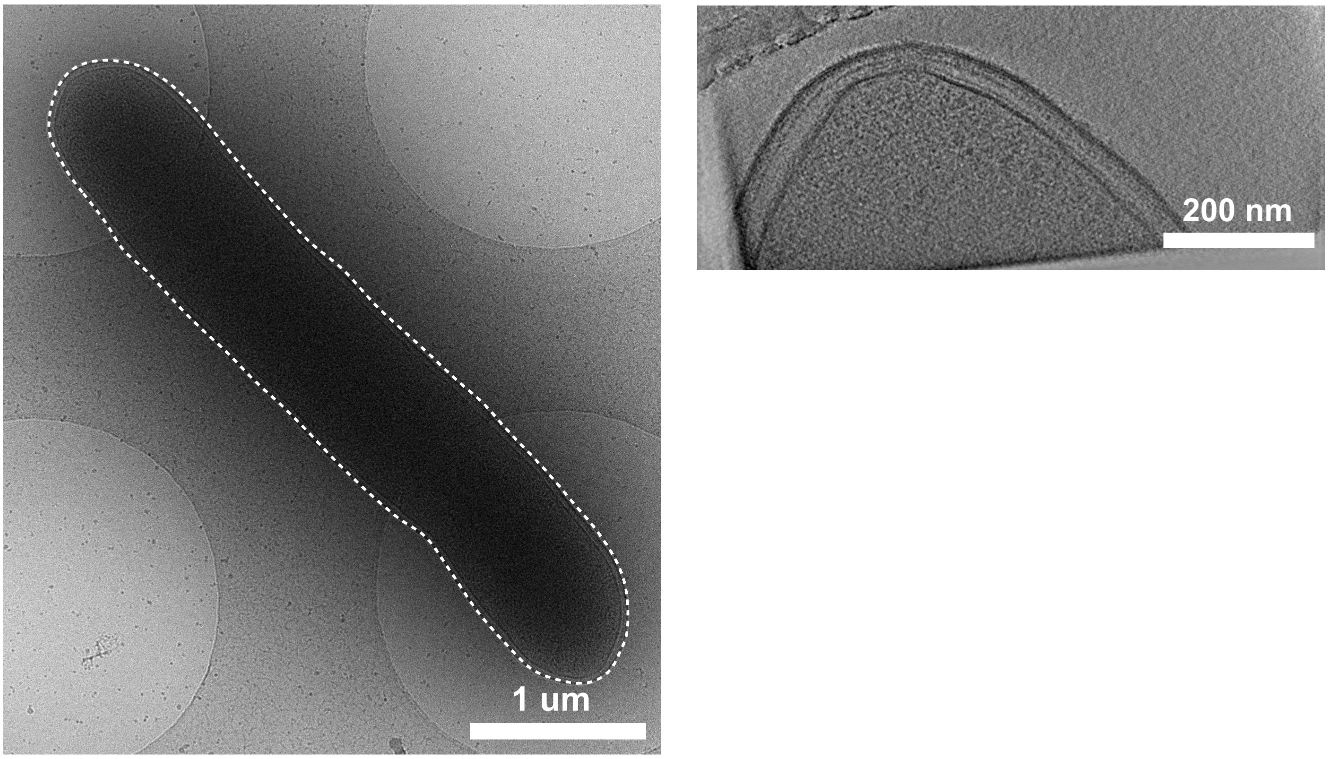


<u>*Bacteroides cellulosyliciticus* DSM 14838</u>

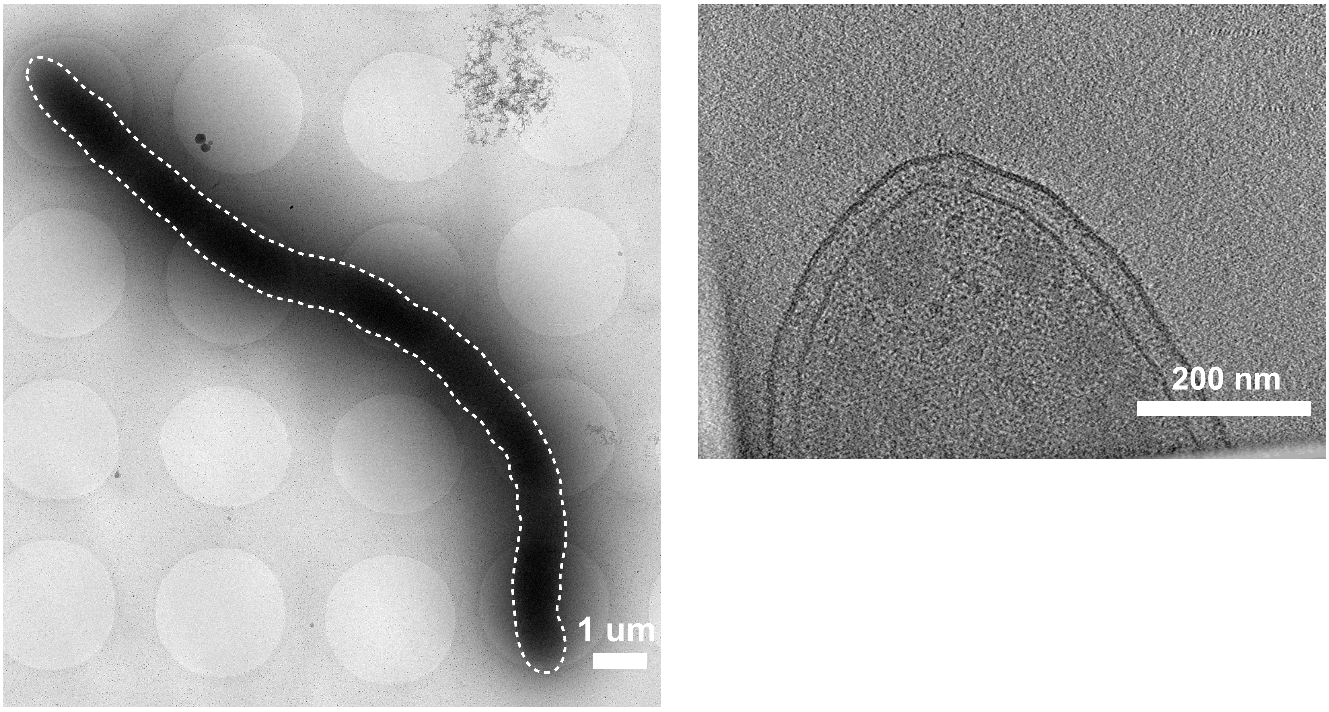

### *Bacteroides thetaiotaomicron* DSM 2079

On Glucose

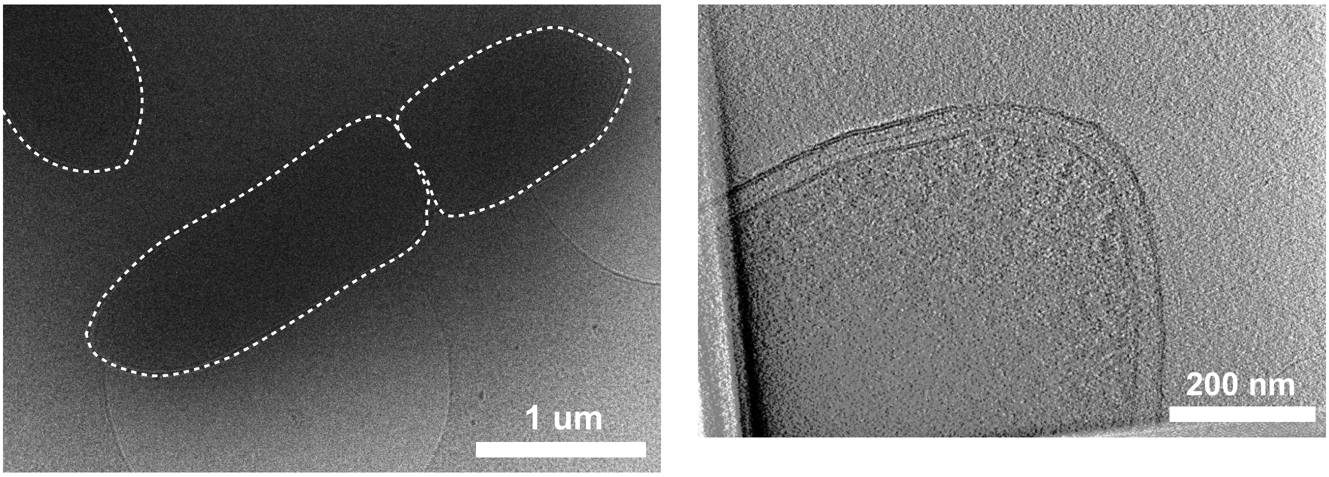


On Starch/Maltose

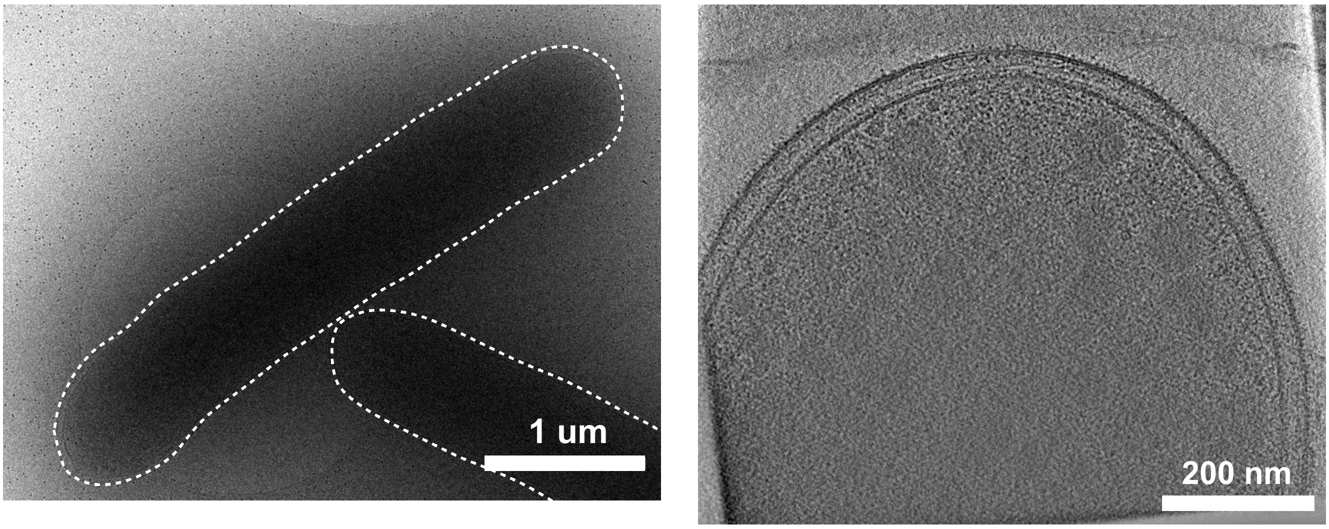


On BHI-Hemin

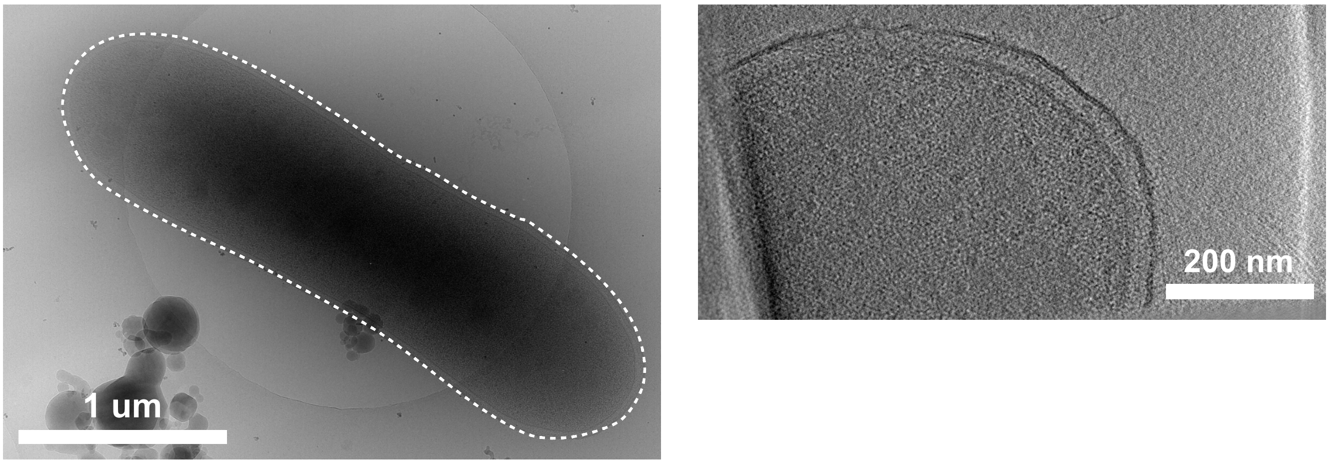

*Bacteroides uniformis* ATCC 8492

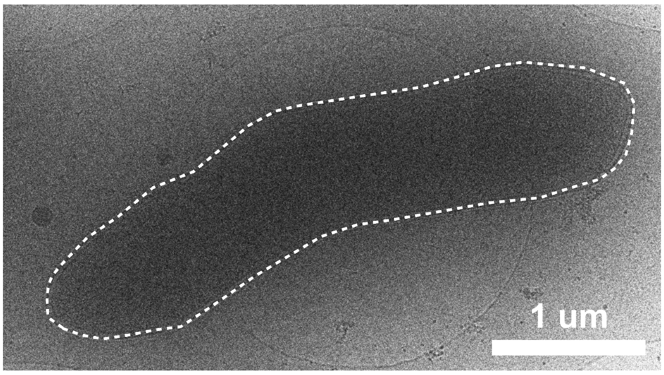


*Bacteroides vulgatus* ATCC 8482

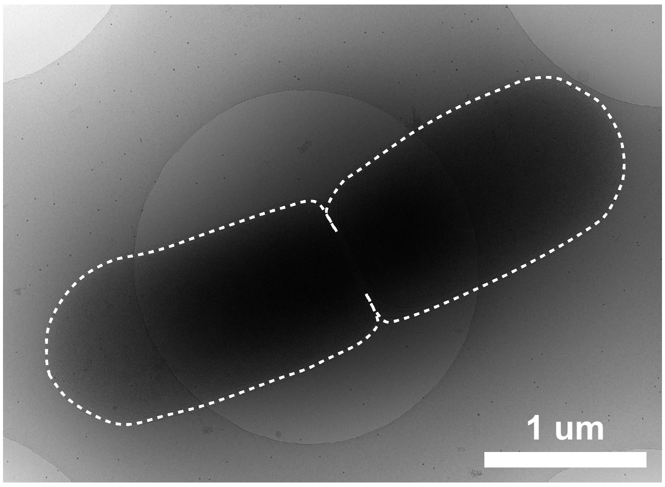


*Basfia succiniproducens* DSM 22022

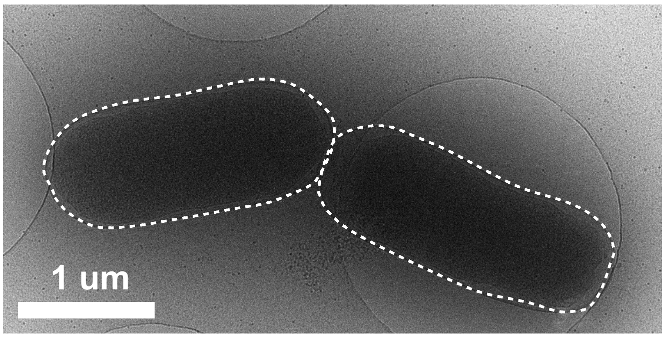


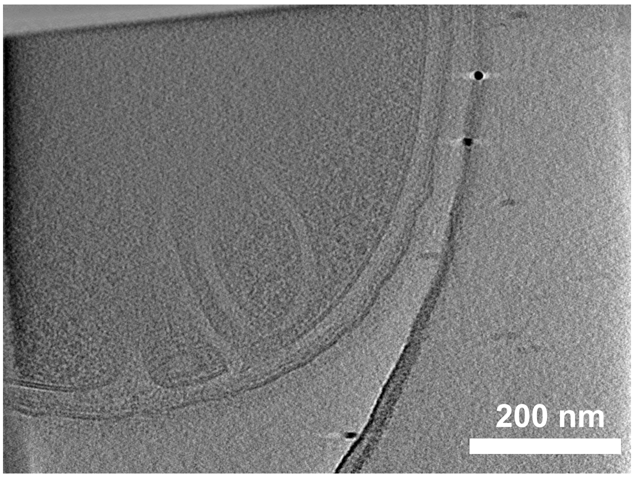


*Bifidobacterium adolescentis* DSM 20087

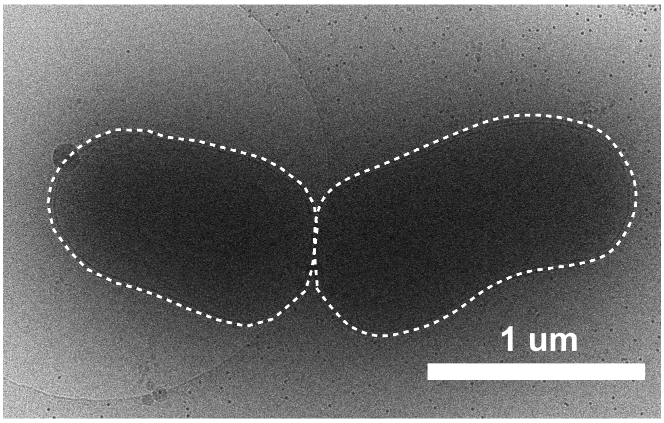


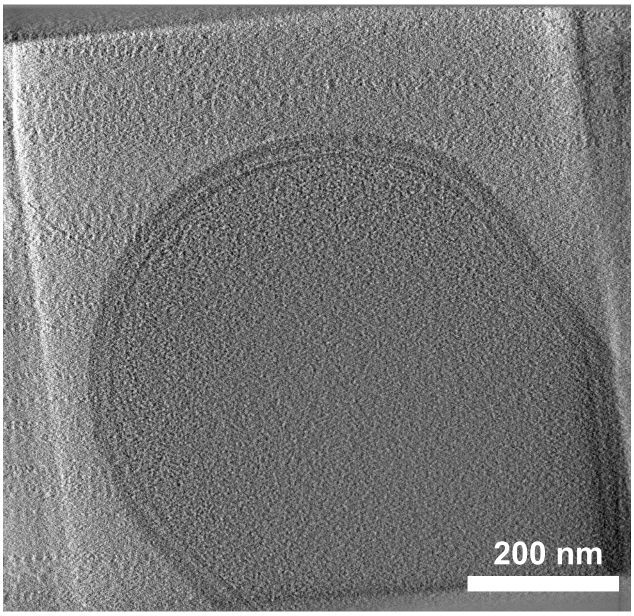

*Bifidobacterium animalis* DSM 10140

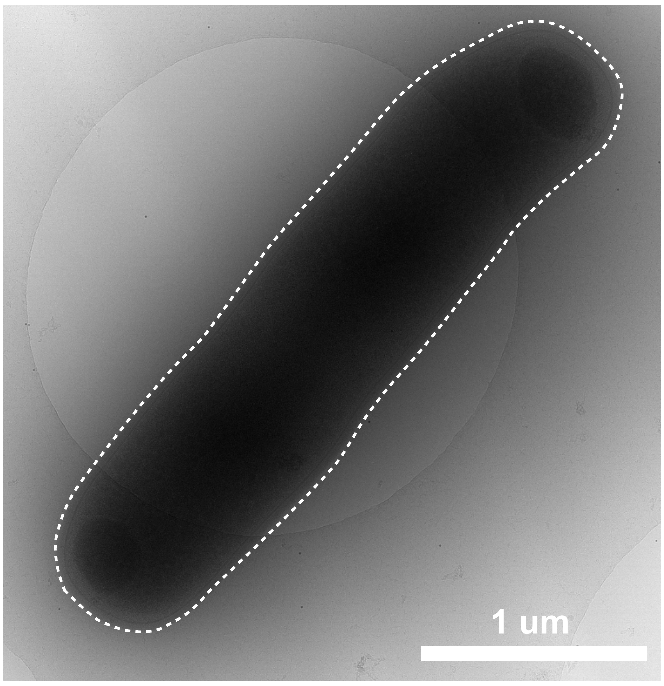


*Bifidobacterium choerinum* AGR 2158

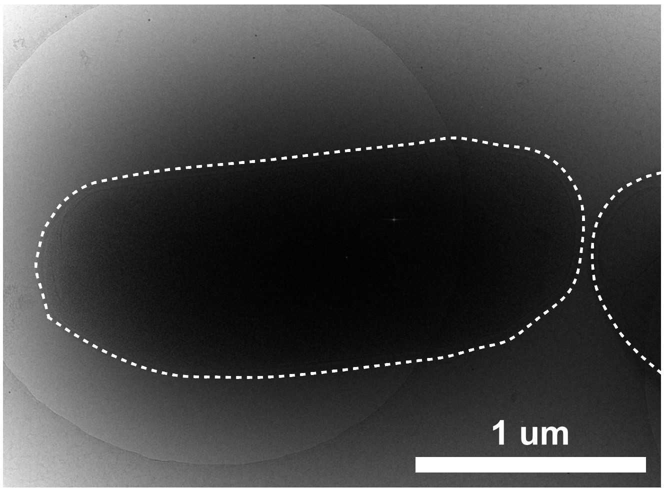


*Bifidobacterium ruminantium* DSM 6489

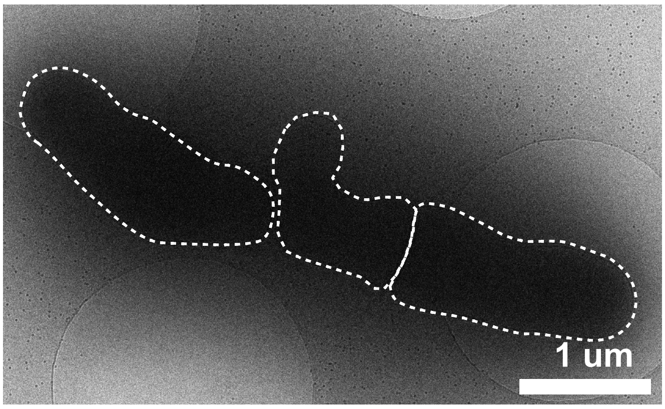


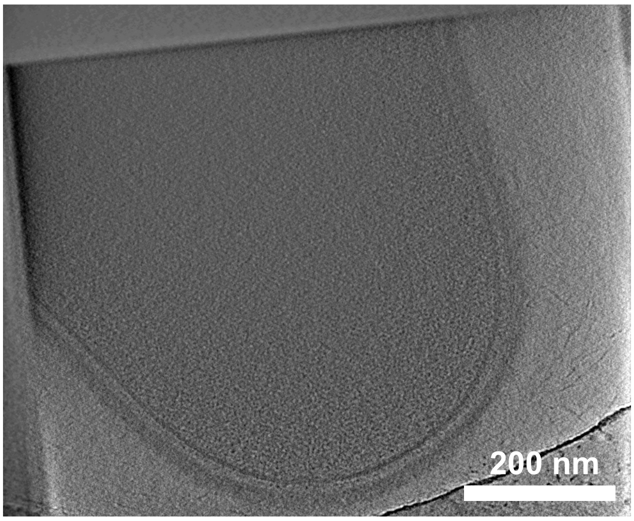


*Bifidobacterium thermophilum* DSM 20209

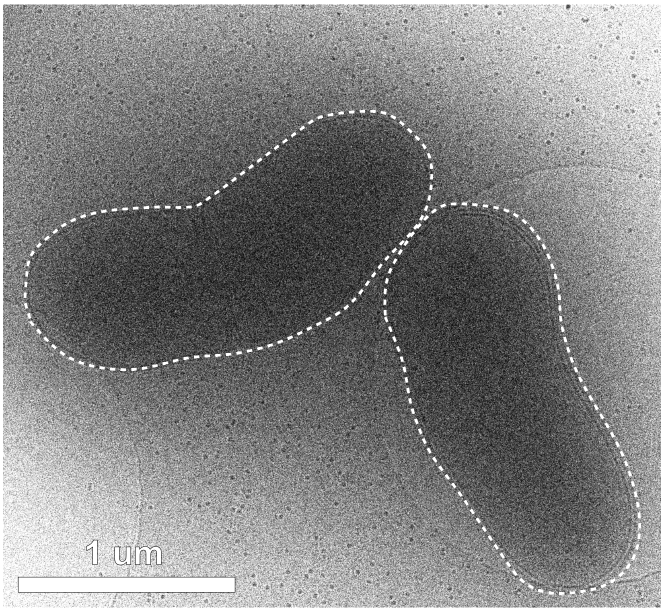


*Blautia schinkii* DSM 10518

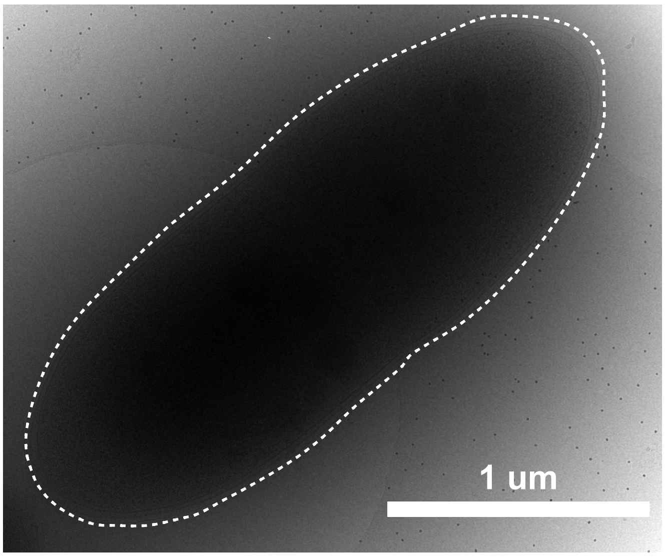

*Butyrivibrio fibrisolvens* CF3

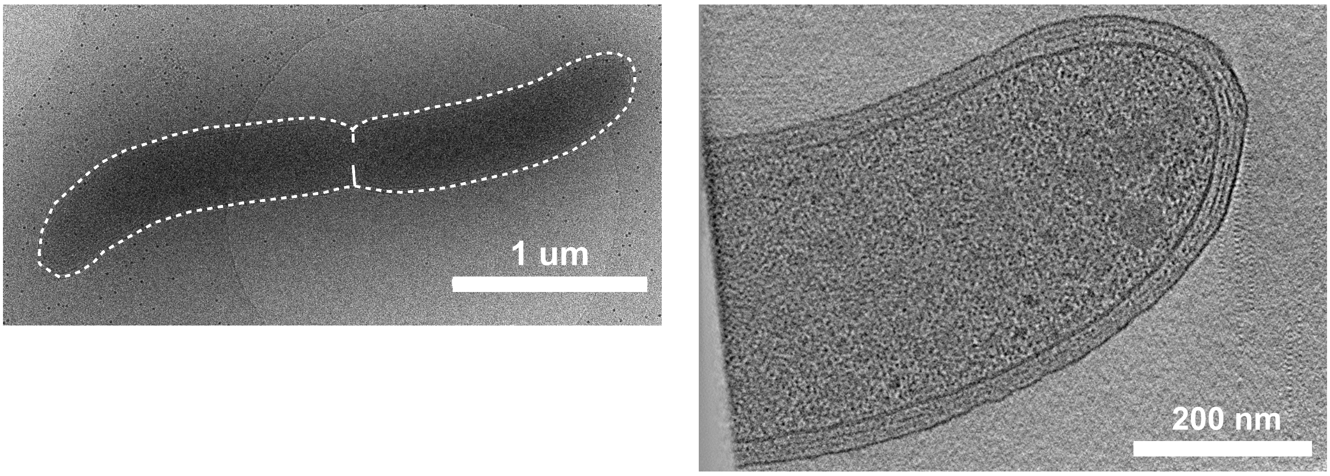


*Butyrivibrio fibrisolvens* D1 DSM 3071

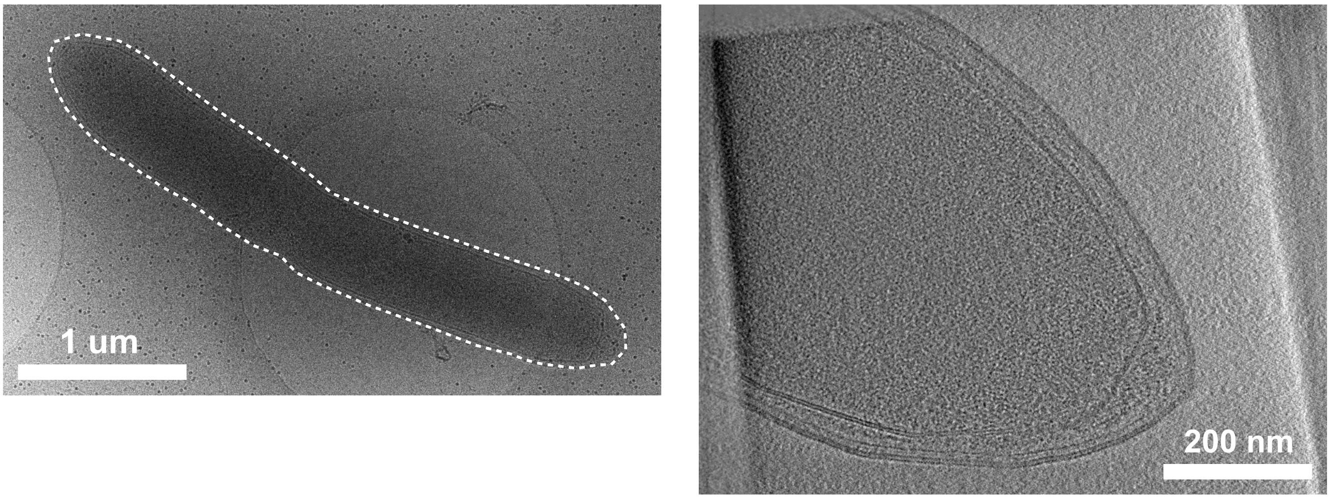


*Butyrivibrio sp.* DSM 10294

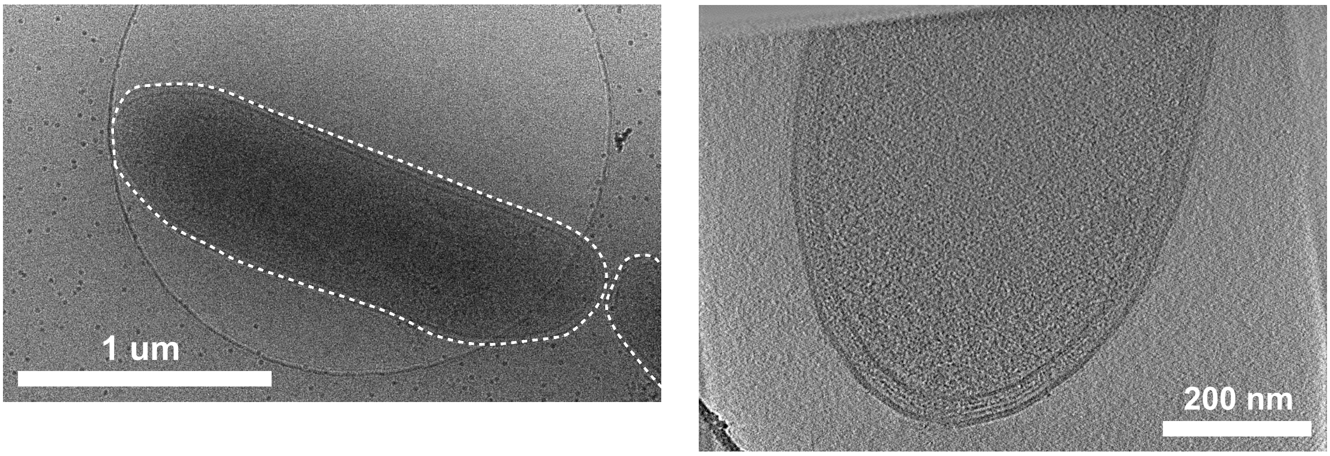

*Clostridium aminophilum* F* DSM 10710

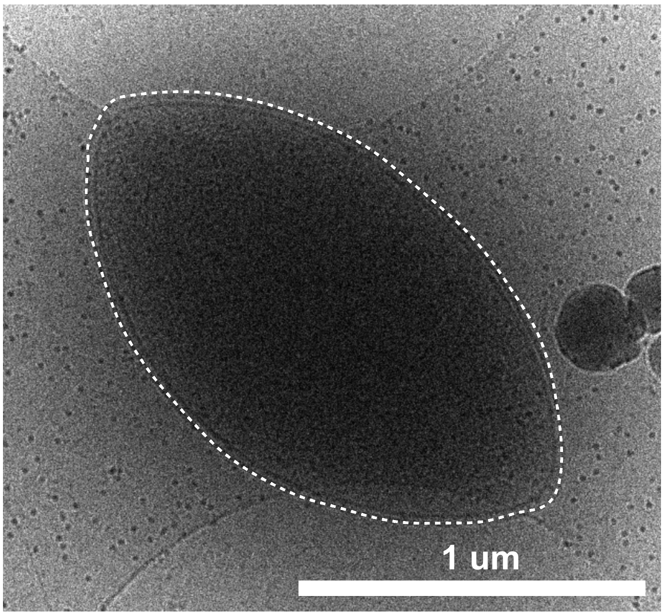


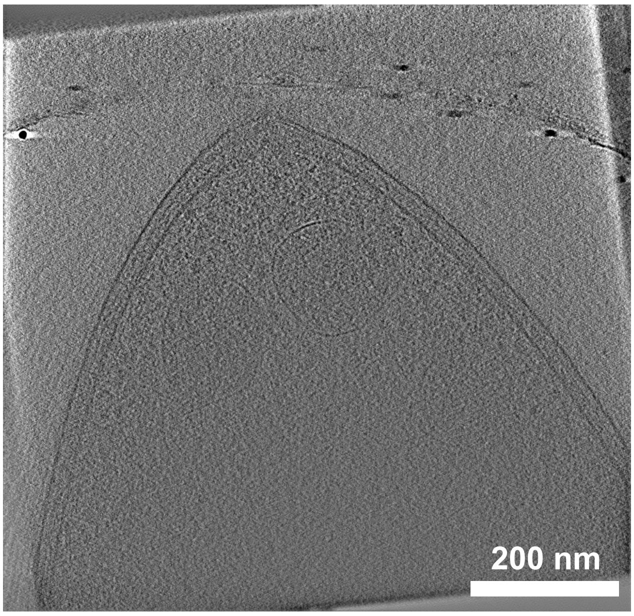


*Clostridium butyricum* 5001 bejerinckii

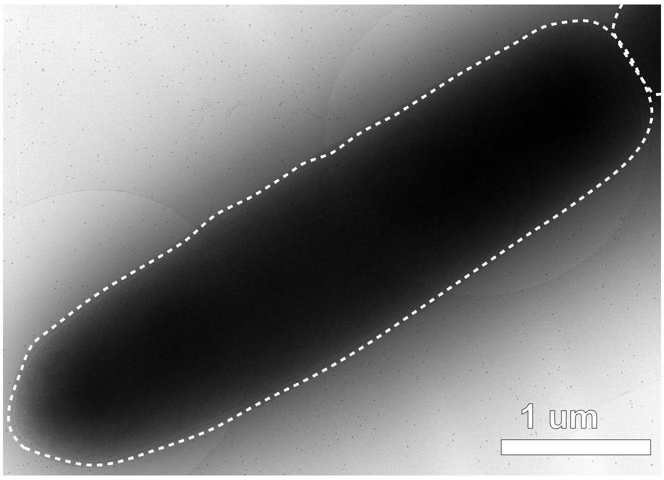


*Clostridium cadaveris* AGR 2141

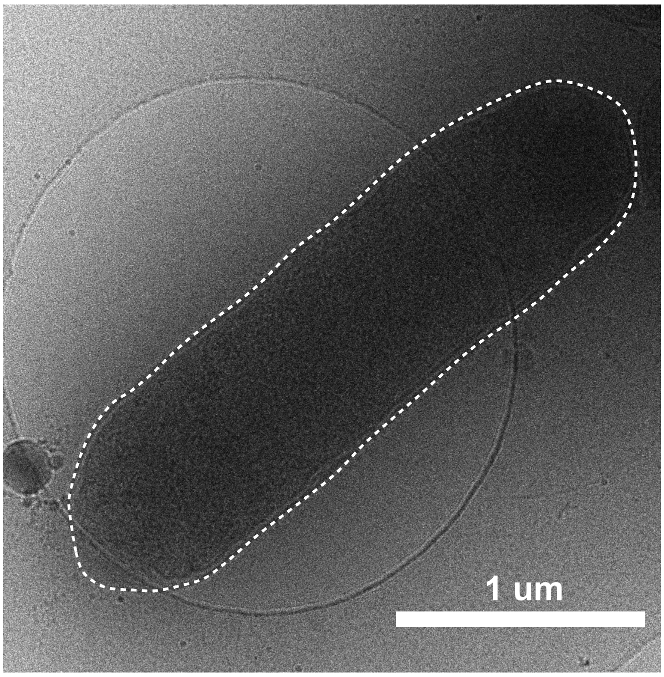


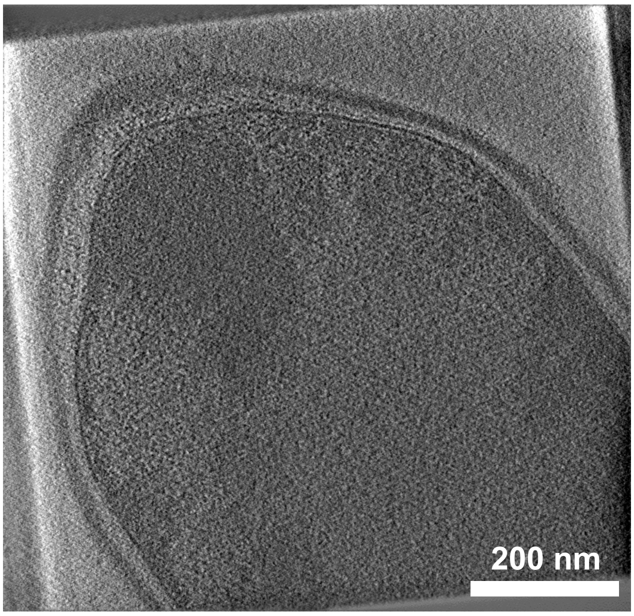

### *Clostridium paraputrificum* AGR 2156

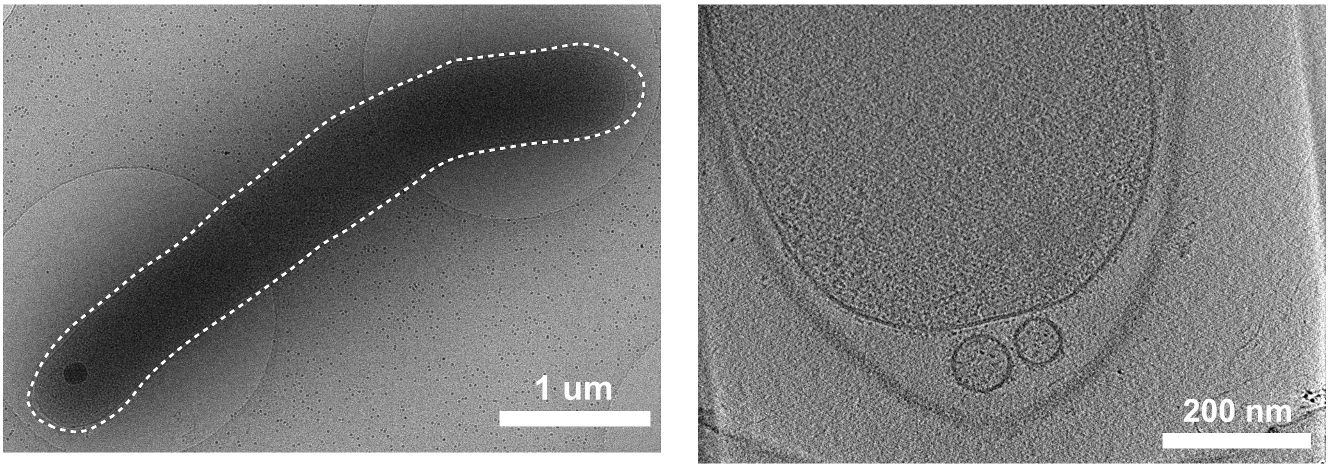


### *Corynebacterium vitaeruminis* Ga6A13

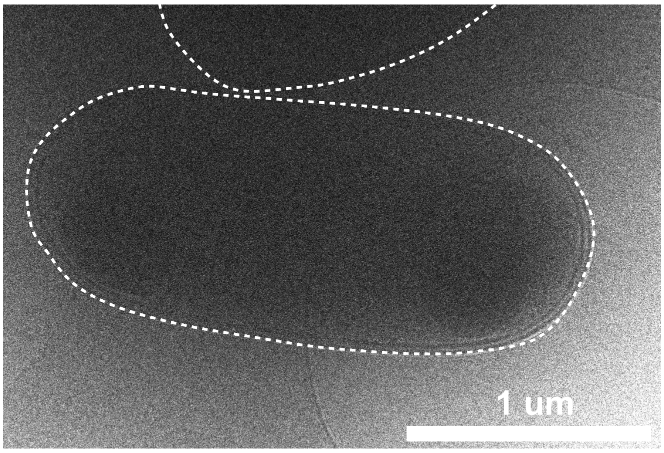


### *Dorea sp.* AGR 2135

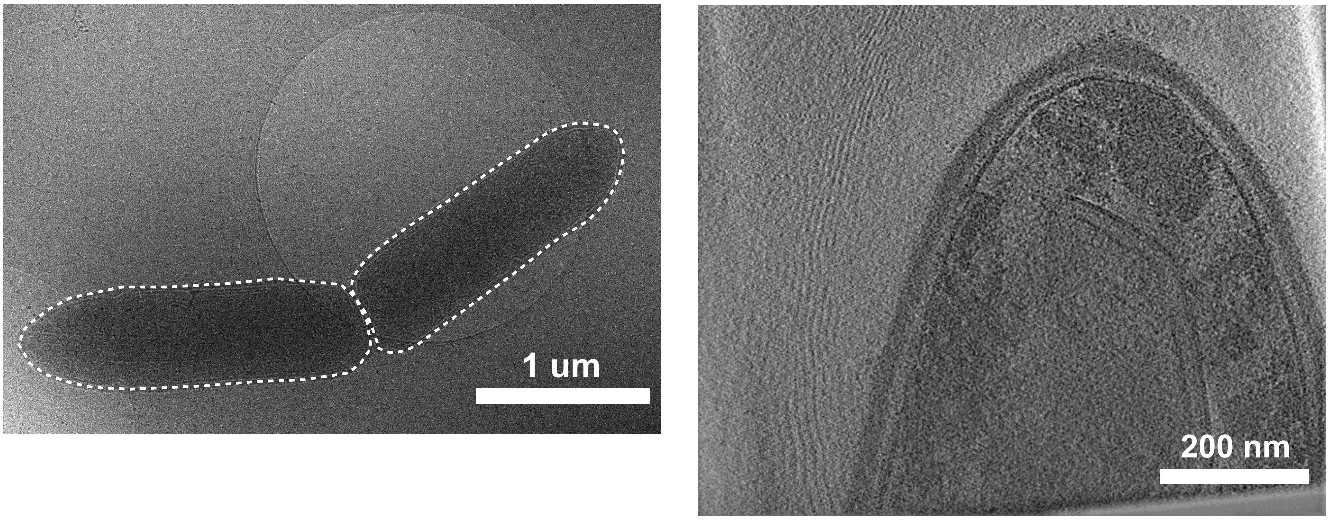


### *Eubacterium pyruvativorans* KHPC4

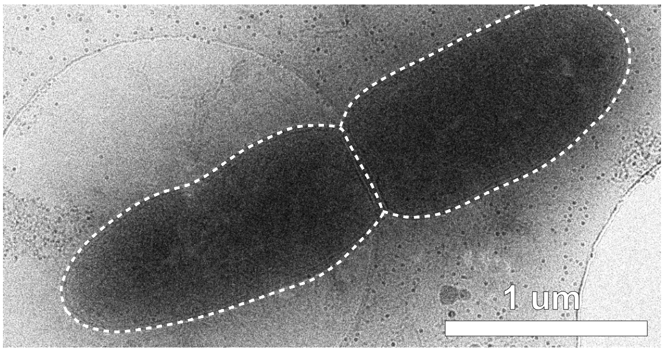


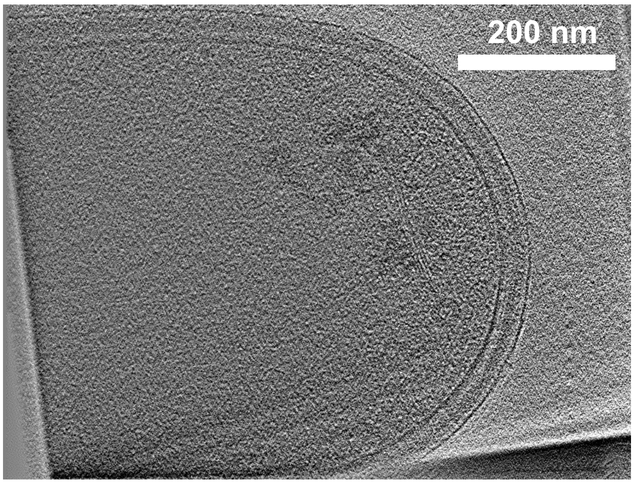

*Eubacterium siraeum* DSM 15702

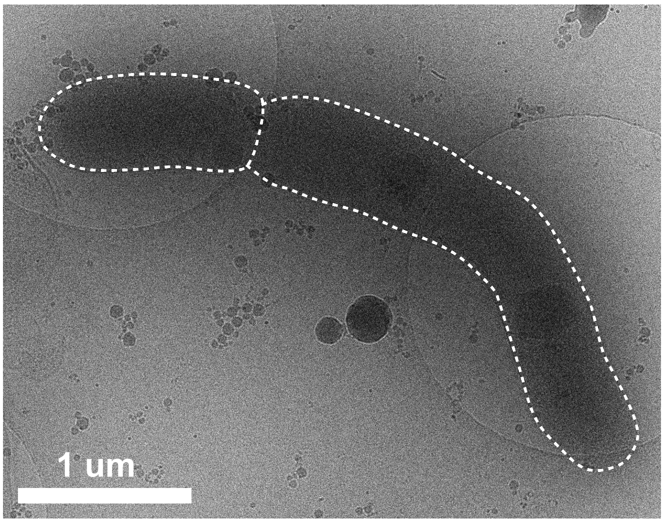


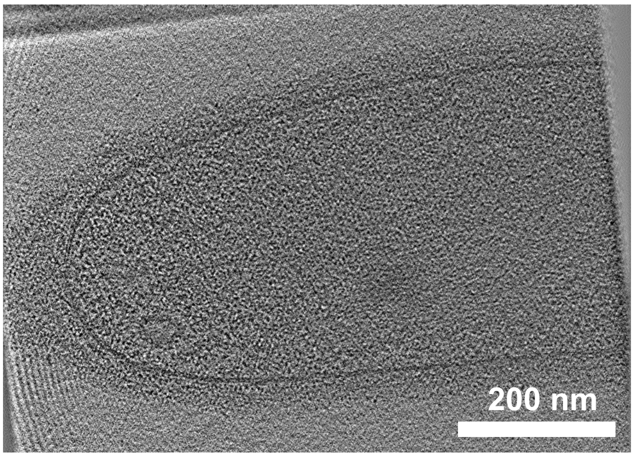


*Faecalibacterium prausnitzii* DSM 17677

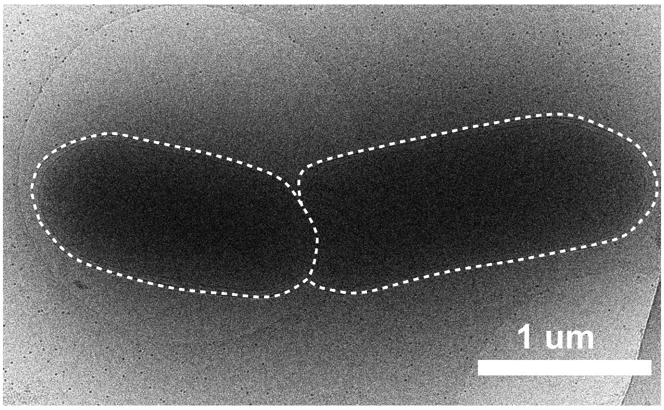


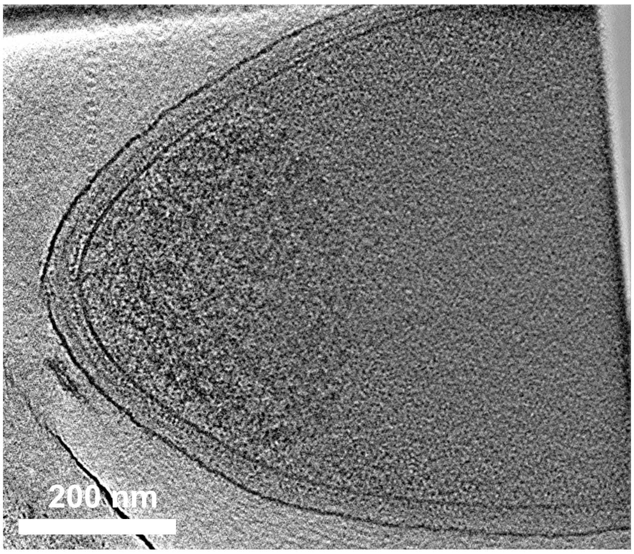


*Fibrobacter succinogenes* ATCC 19169

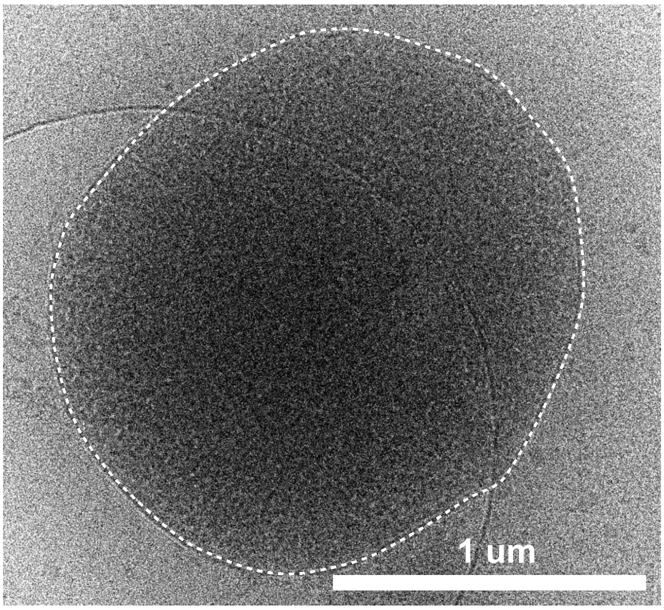


*Fusobacterium necrophorum* Hun 048

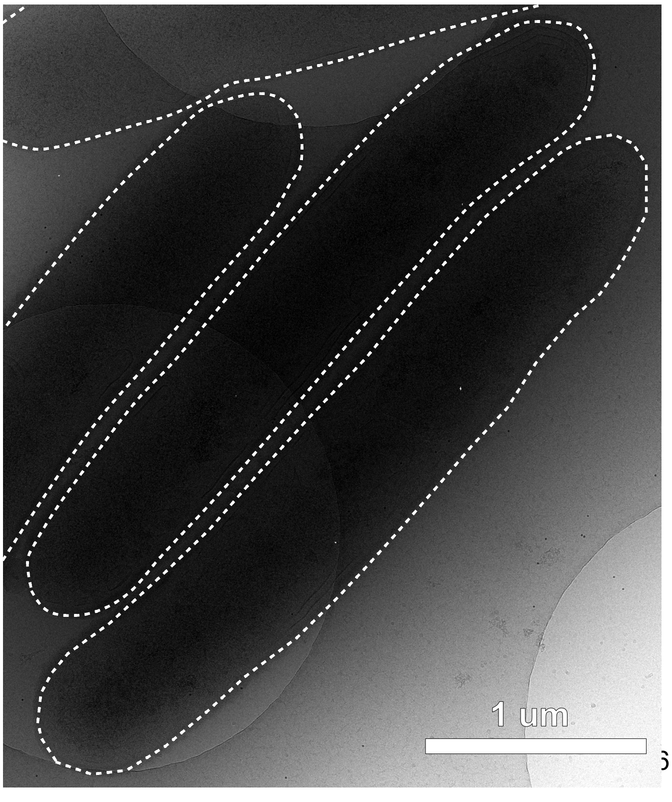

*Kandleria vitulina* DSM 20405

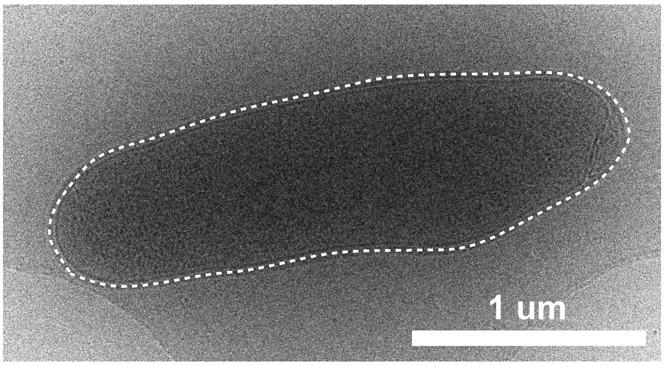


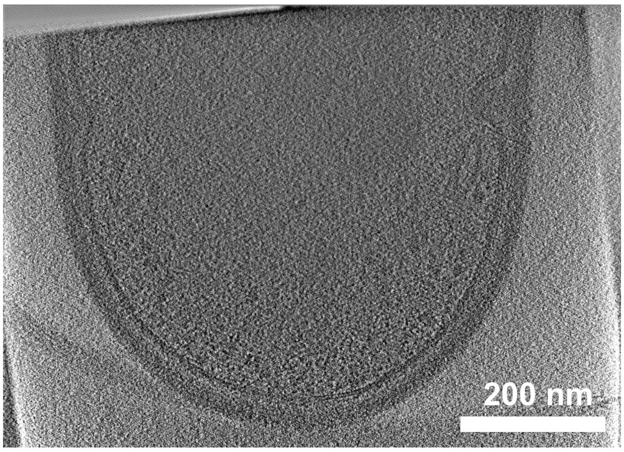


*Lachnobacterium bovis* DSM 14045

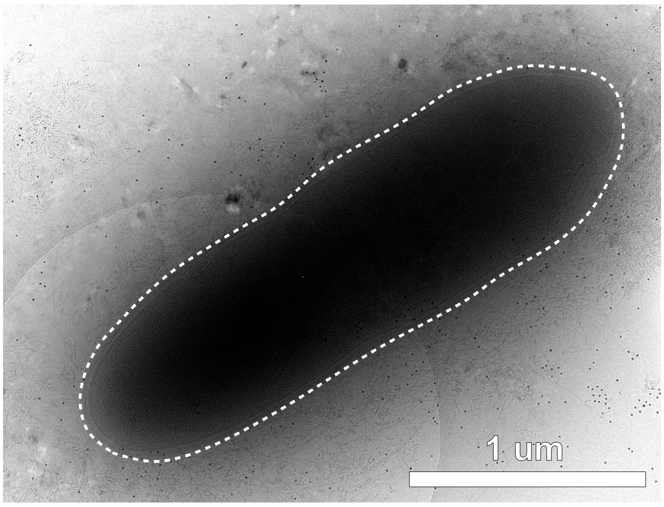


*Lachnospira multipara* ATCC 19207

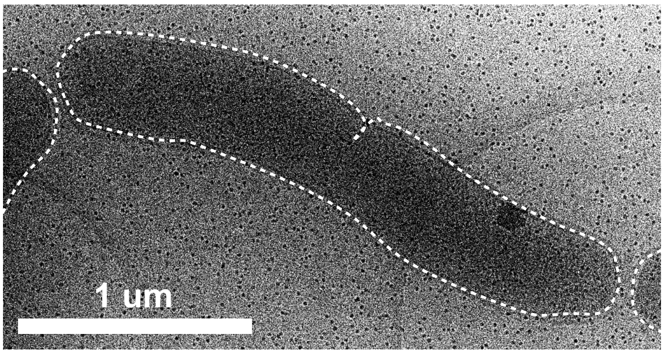


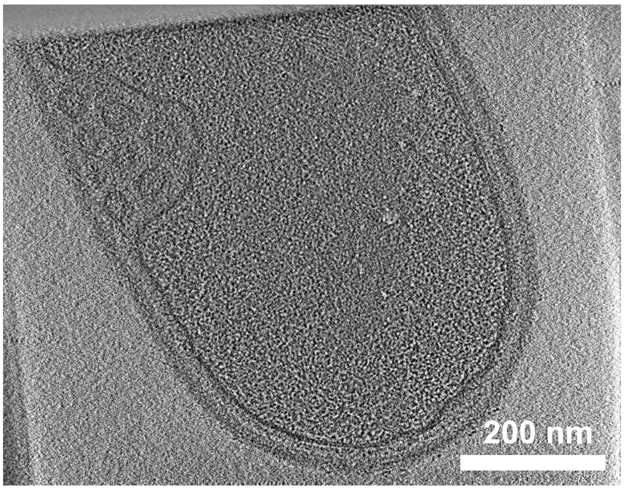


*Ligilactobacillus ruminis* DSM 20403

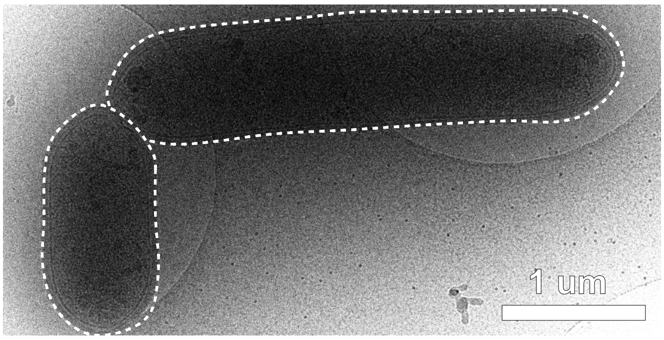


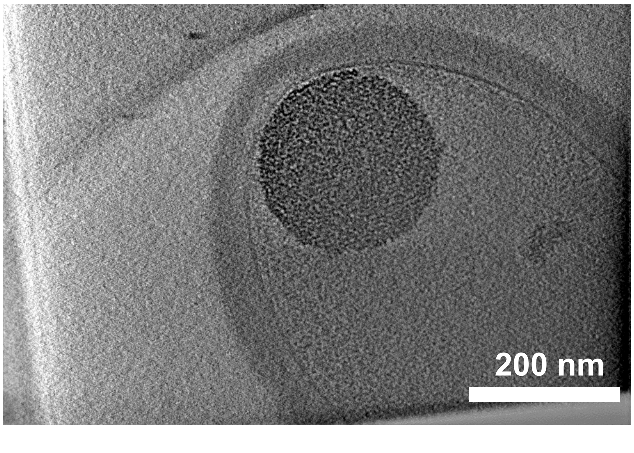

*Limosilactobacillus mucosae* AGR 63

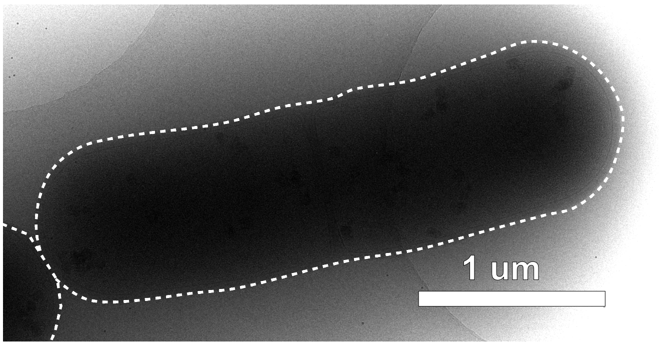


*Limosilactobacillus reuteri* DSM 17938

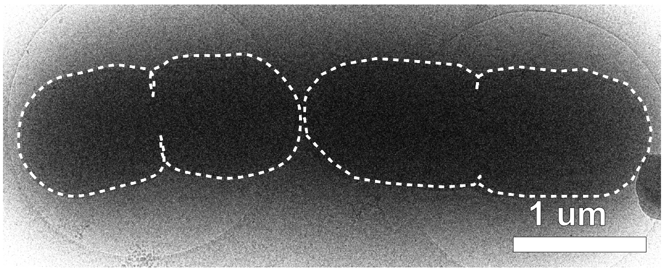


*Marvinbryantia formatexigens* DSM 14469

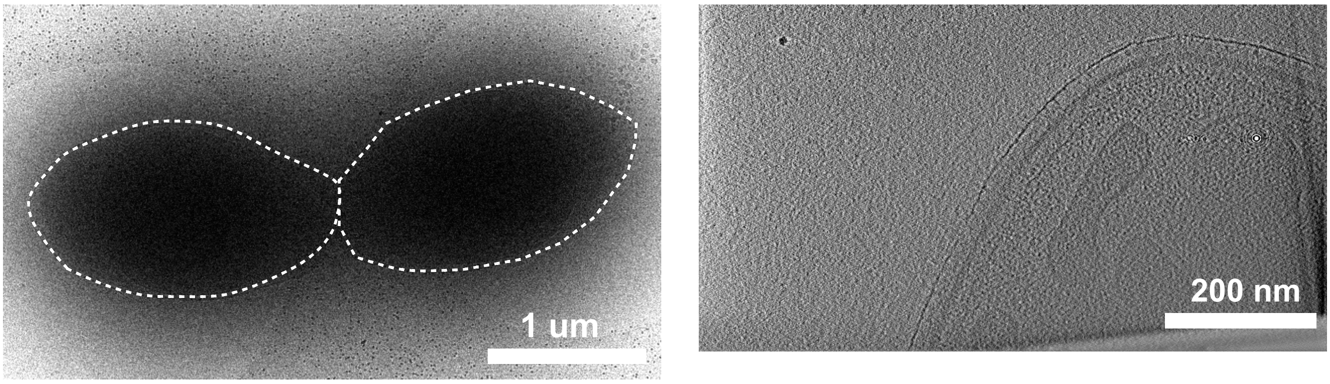


*Megamonas sp.* Calf 98-2

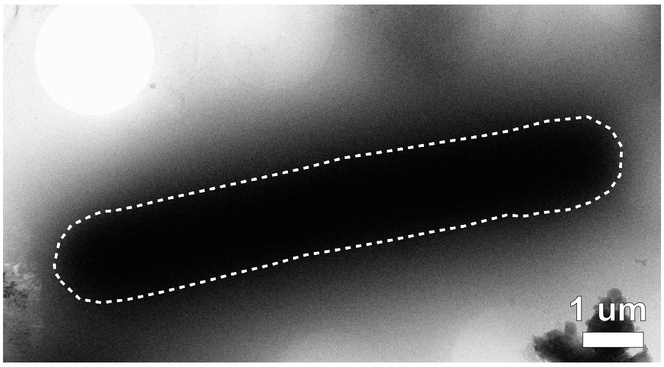


*Megasphaera elsdenii* DSM 20460

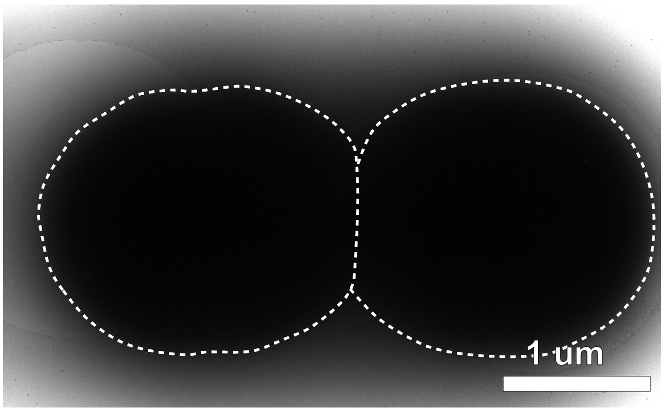


*Methanobrevibacter ruminantium* DSM 1093

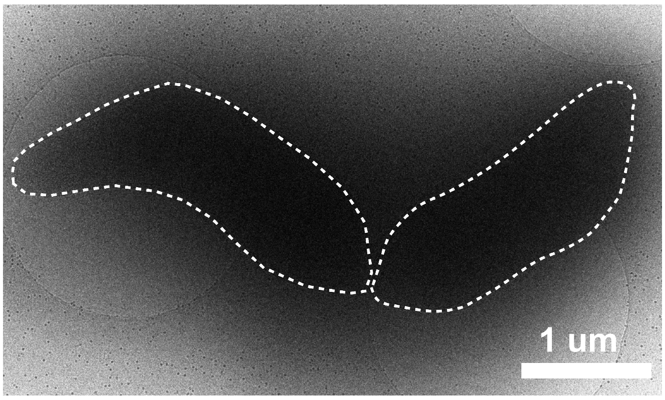

*Methanocorpusculum labreanum* DSM 4855

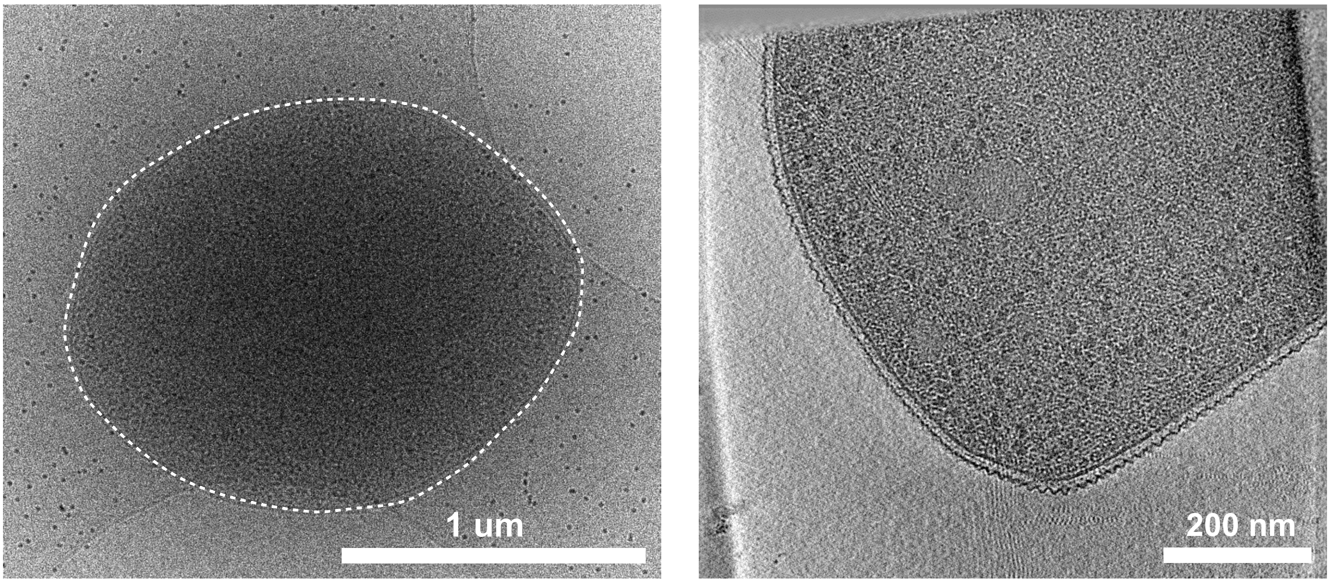


*Methanomicrobium mobile* DSM 1539

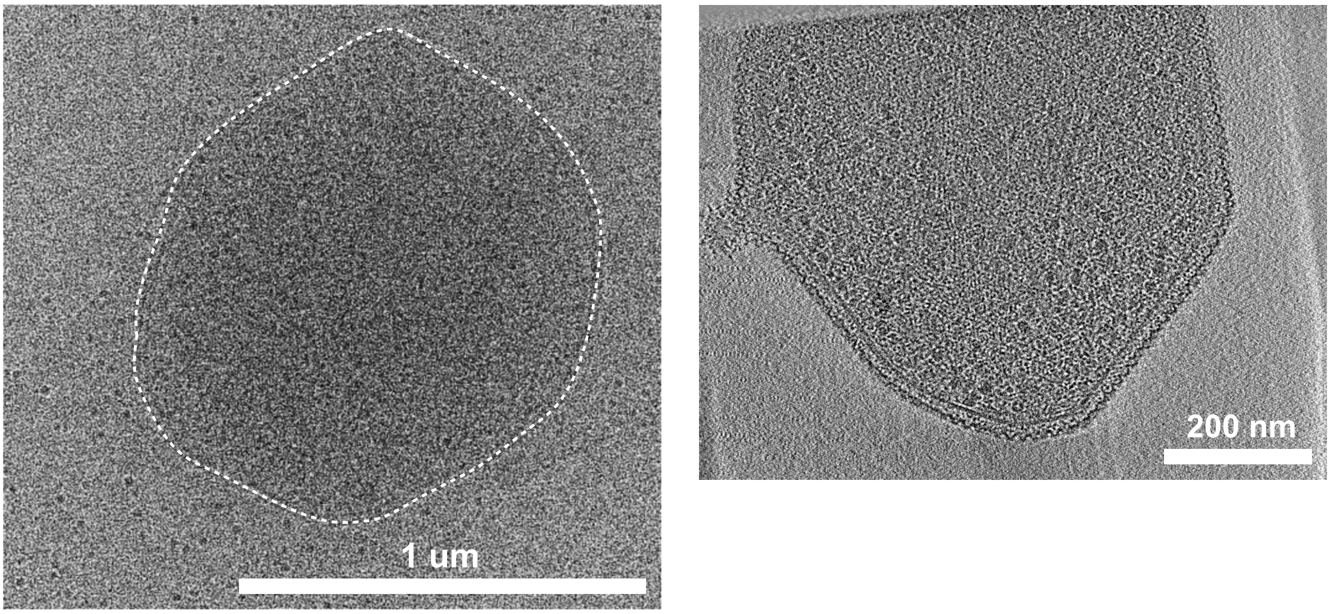


*Micrococcales bacterium* KH10

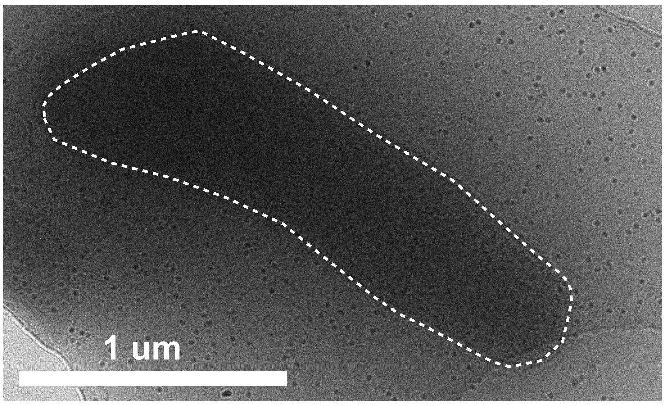


*Mitsuokella jalaludinii* DSM 13811

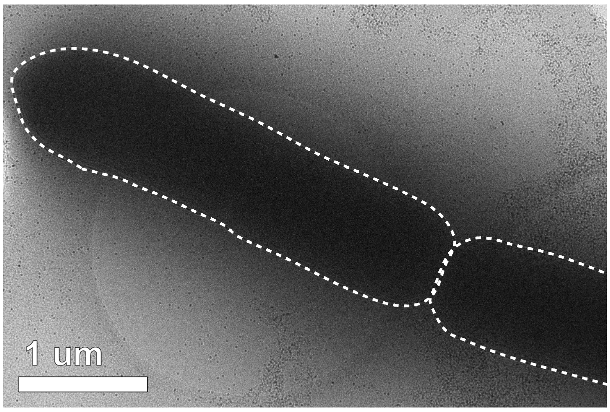

*Olsenella umbonata* WCP15

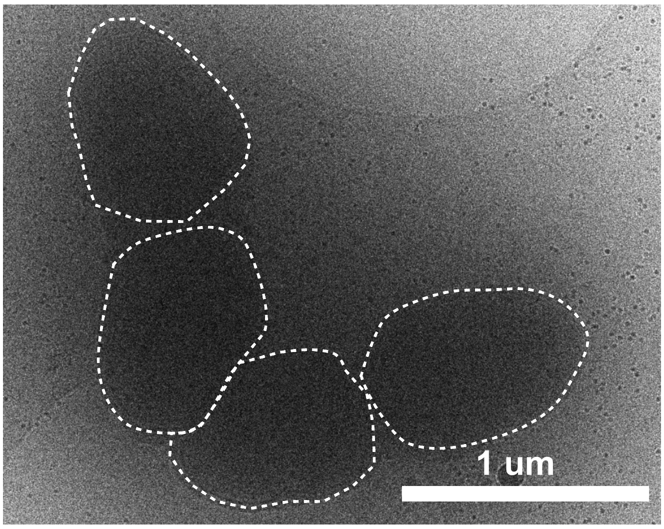


*Pediococcus acidilactici* AGR 20

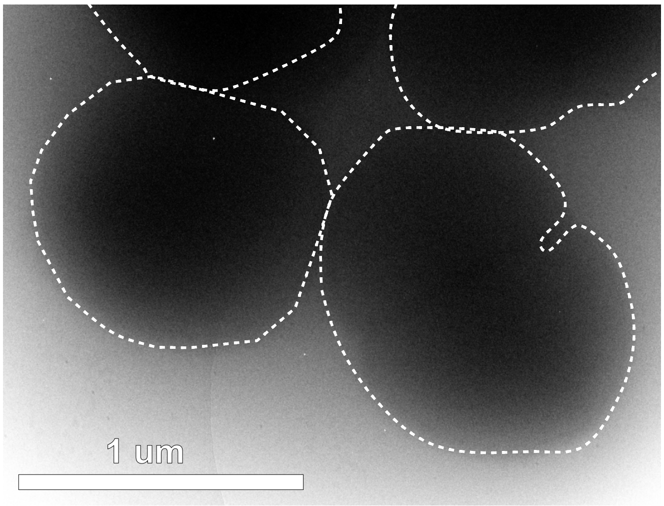


*Peptostreptococcus anaerobius* C

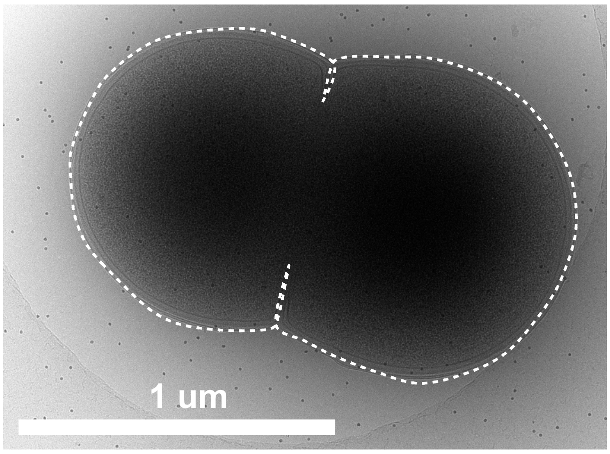


*Prevotella bryantii* DSM 11371

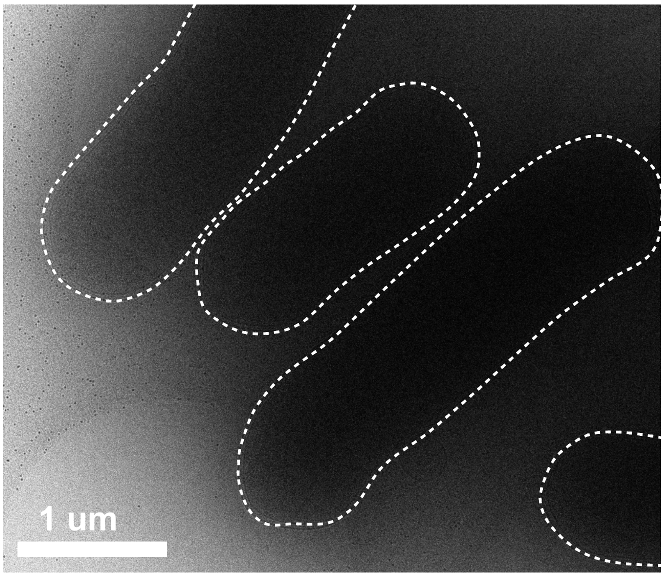


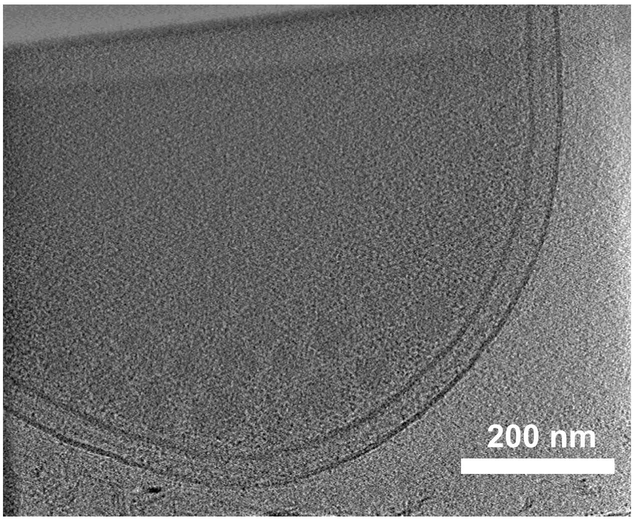

*Prevotella ruminicola* ATCC 19189

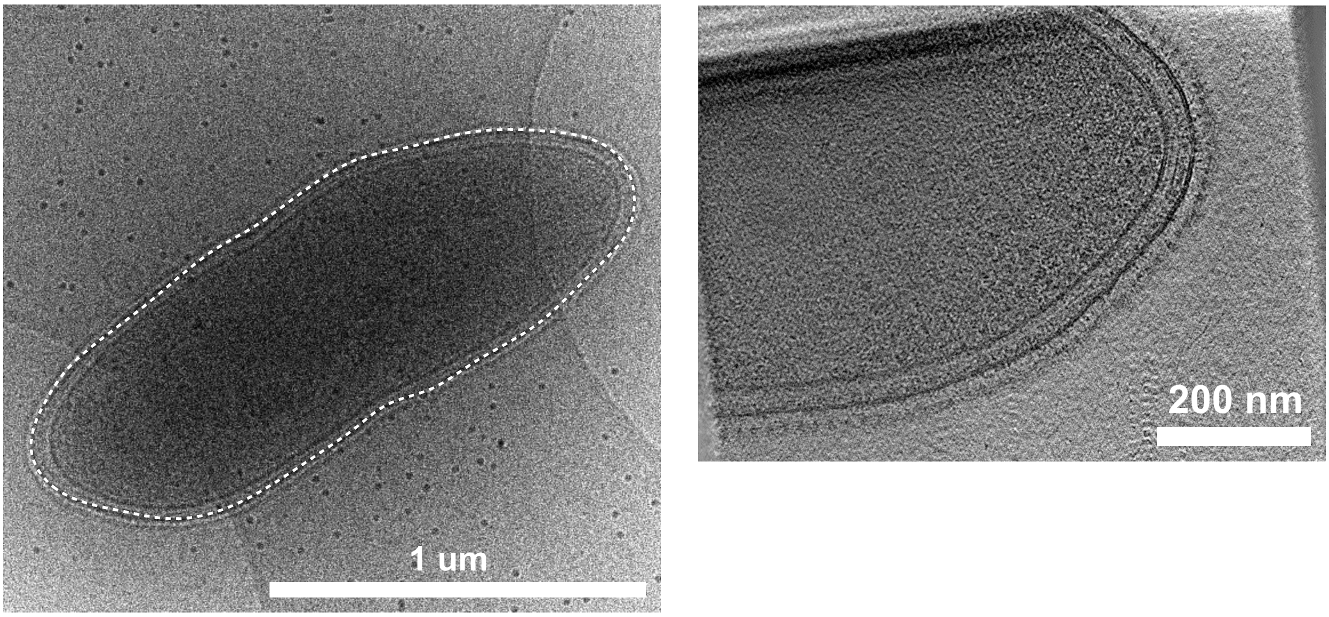


*Propionibacteriaceae bacterium* P6A17

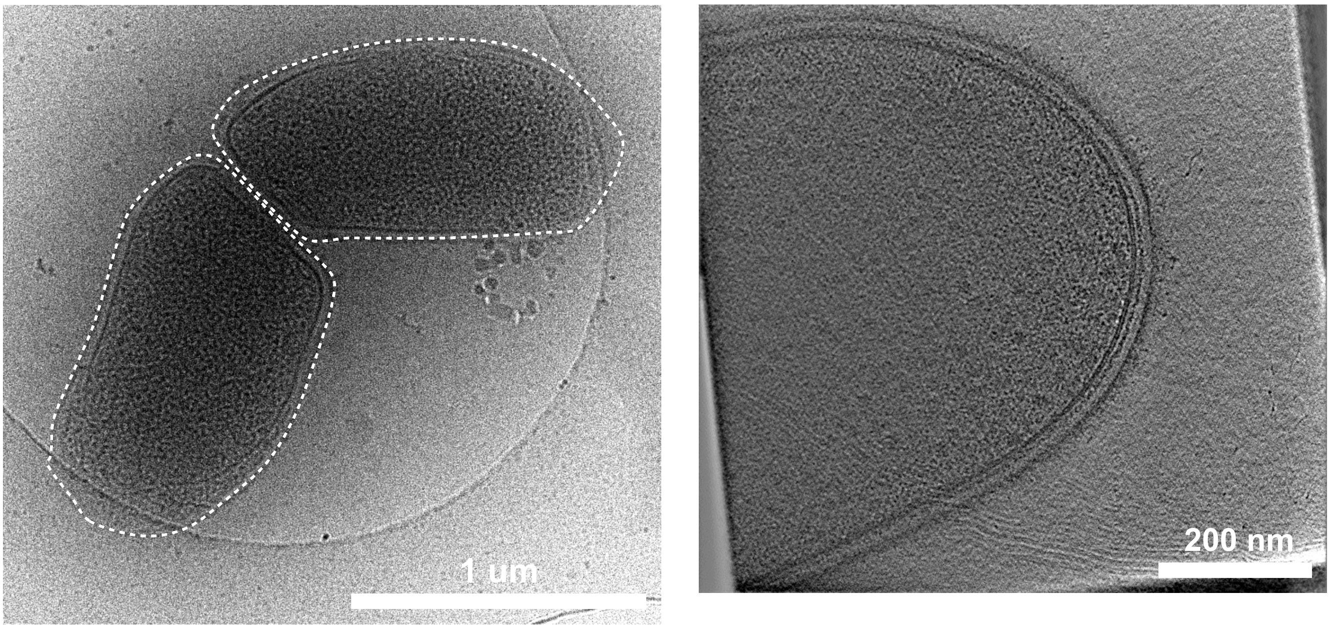


*Proteiniclasticum ruminis* DSM 24773

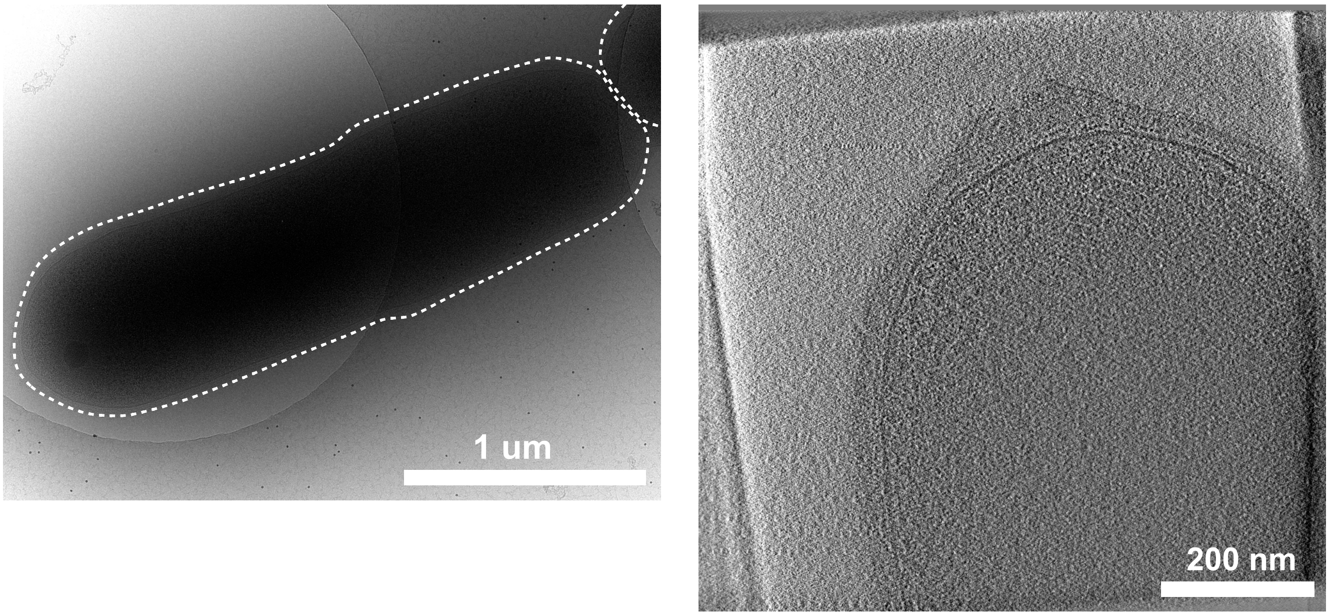

*Pseudobutyrivibrio sp.* LB2011

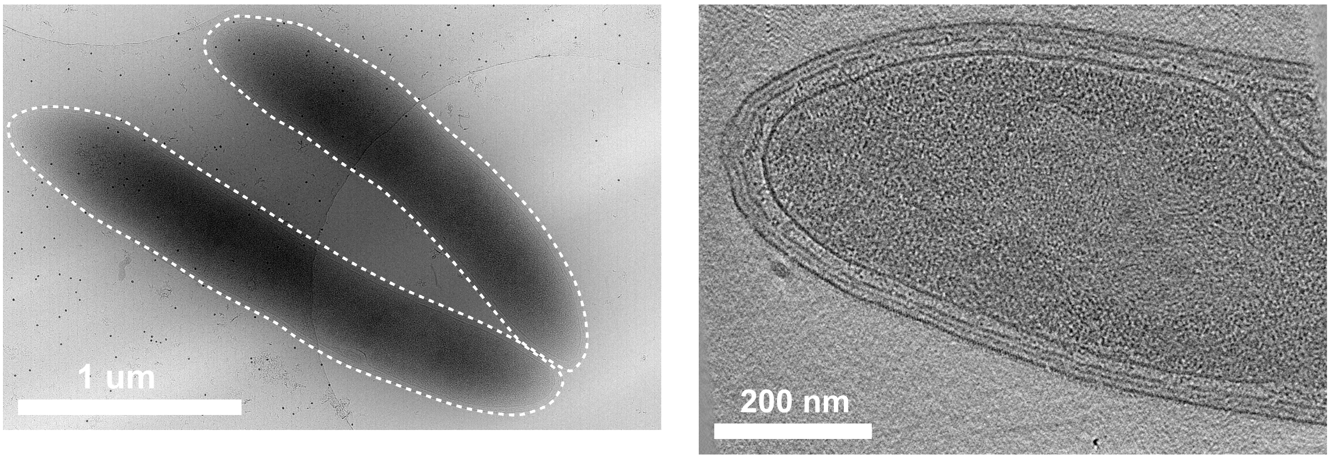


*Roseburia intestinalis* DSM 14610

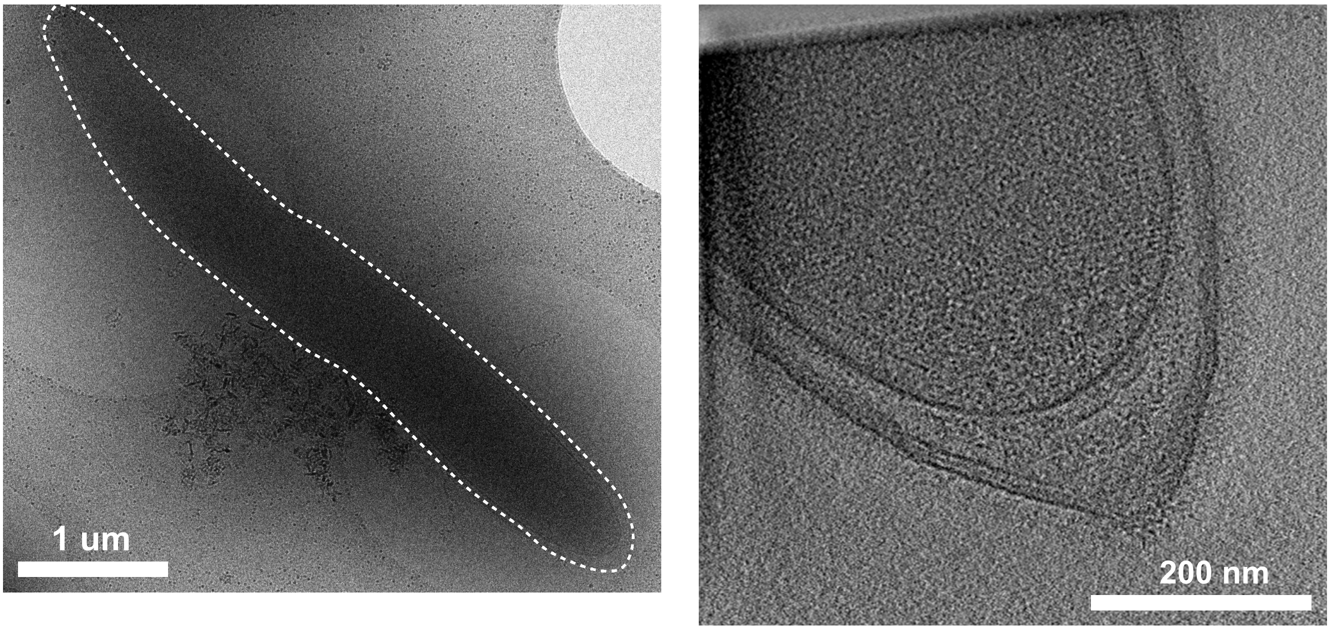


*Roseburia inolinivorans* DSM 16841

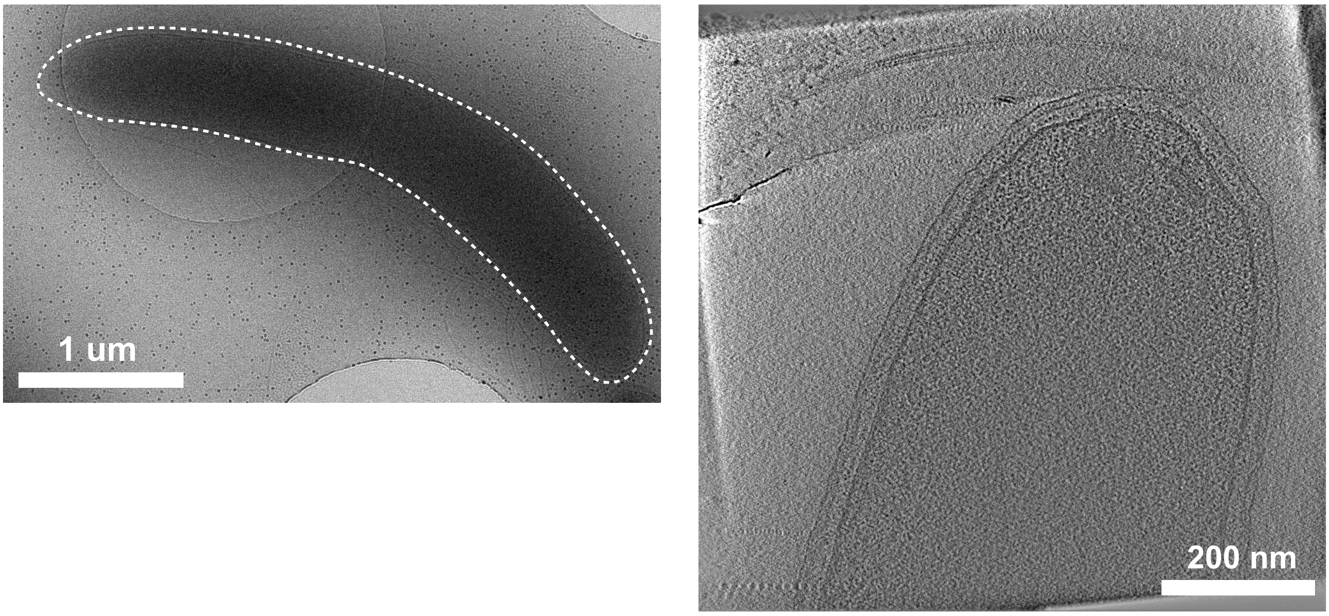

### *Ruminococcus albus* DSM 20455

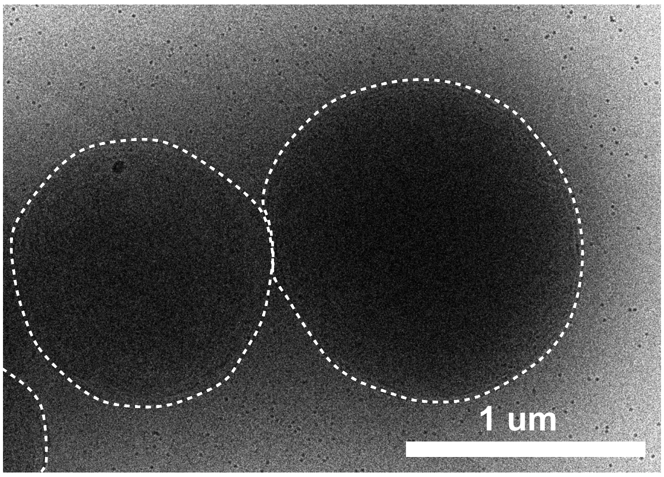


### *Ruminococcus bromii* L2-63

On Fructose

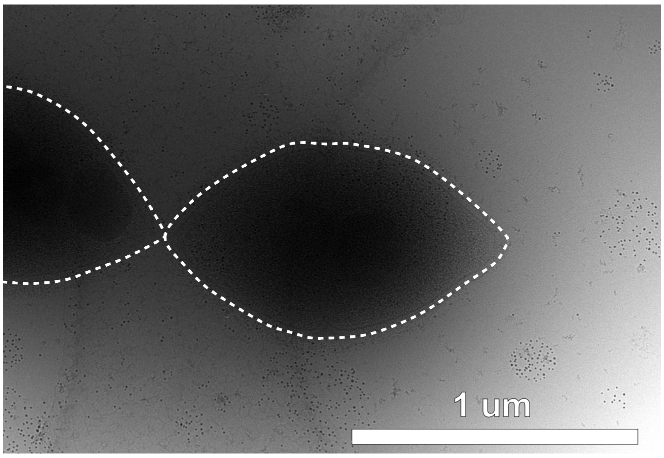


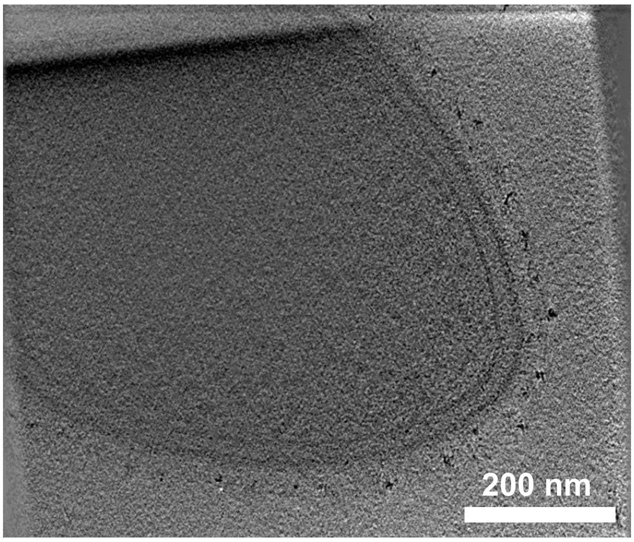


On Pullulan

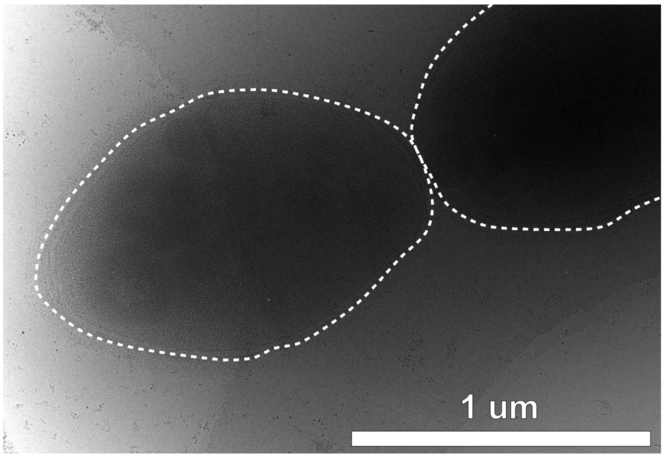


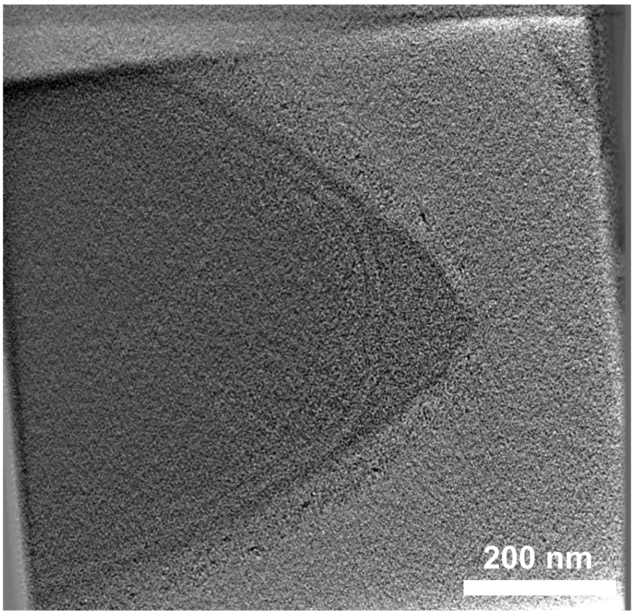

On Starch

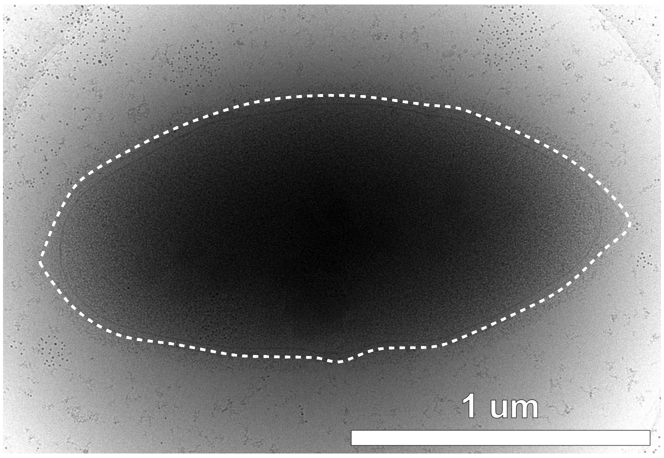


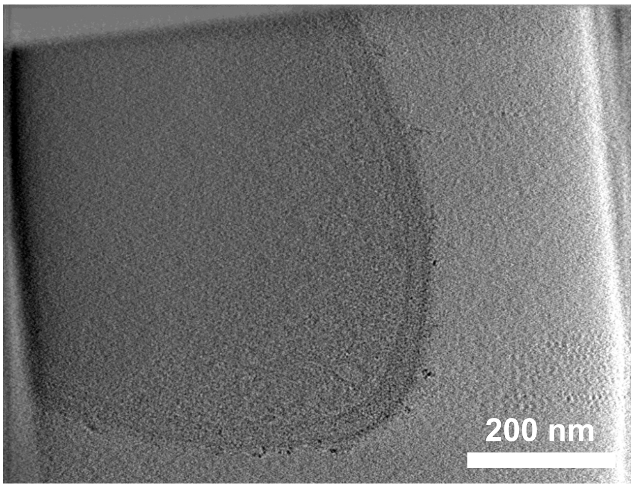


*Ruminococcus champanellensis* JCM 17042

On Cellobiose

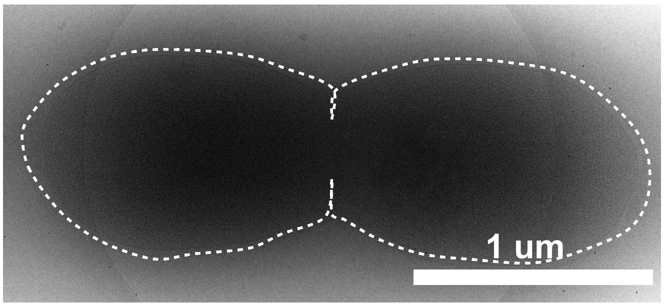


On MCC

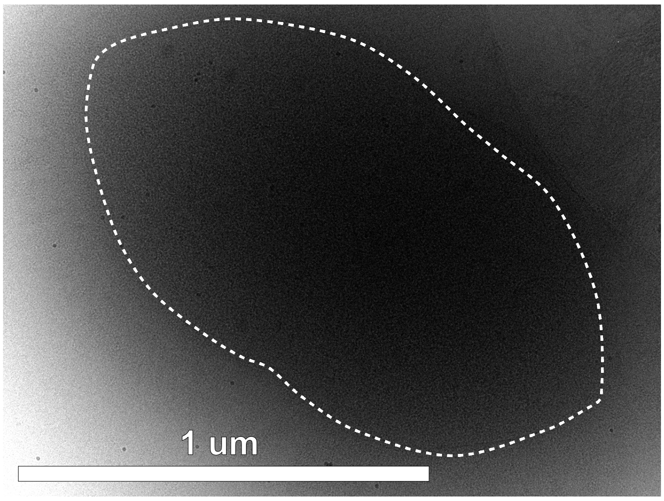


*Ruminococcus gnavus* AGR 2154 / Hun 22

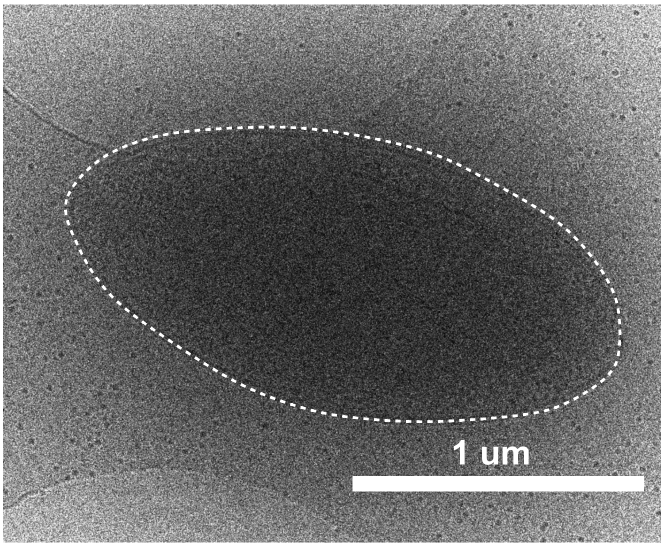

### *Selenomonas bovis* DSM 23594

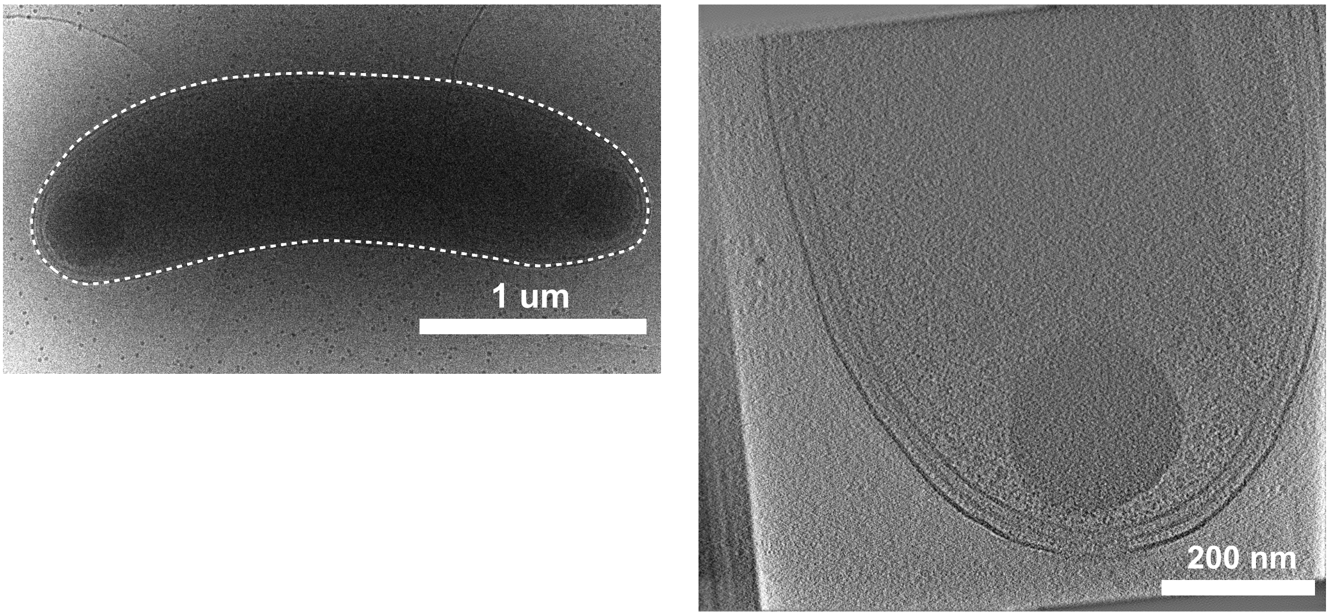


### *Selenomonas ruminantium* AB3002 P159

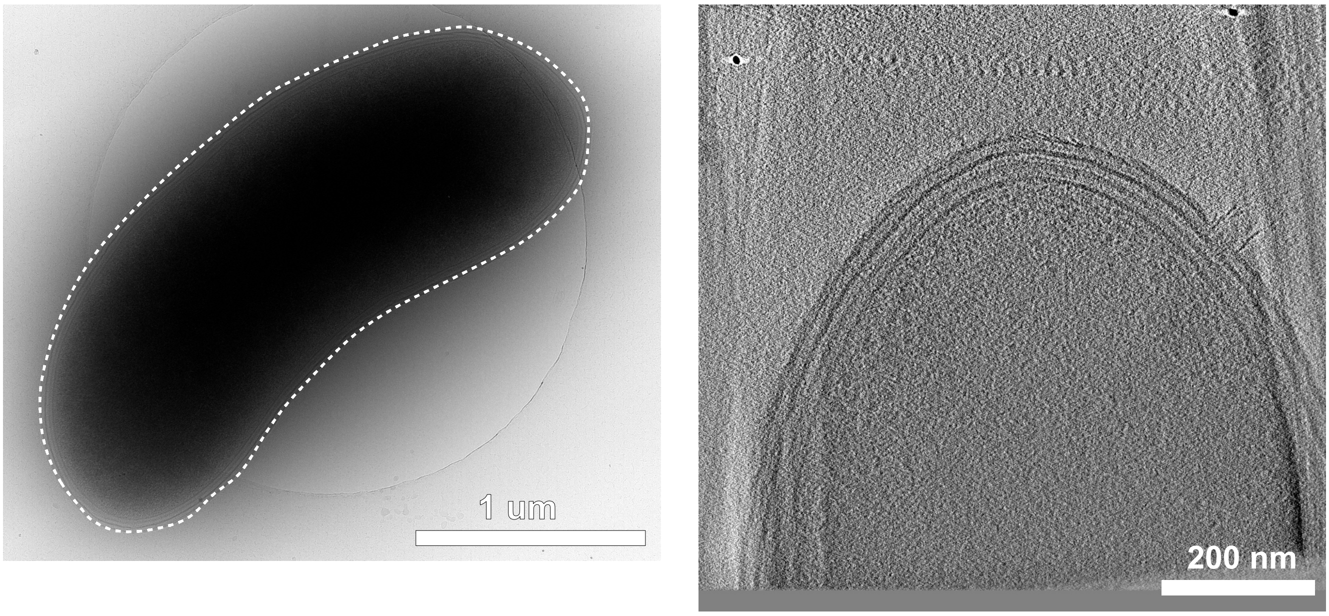


### *Sharpea azabuensis* DSM 20406

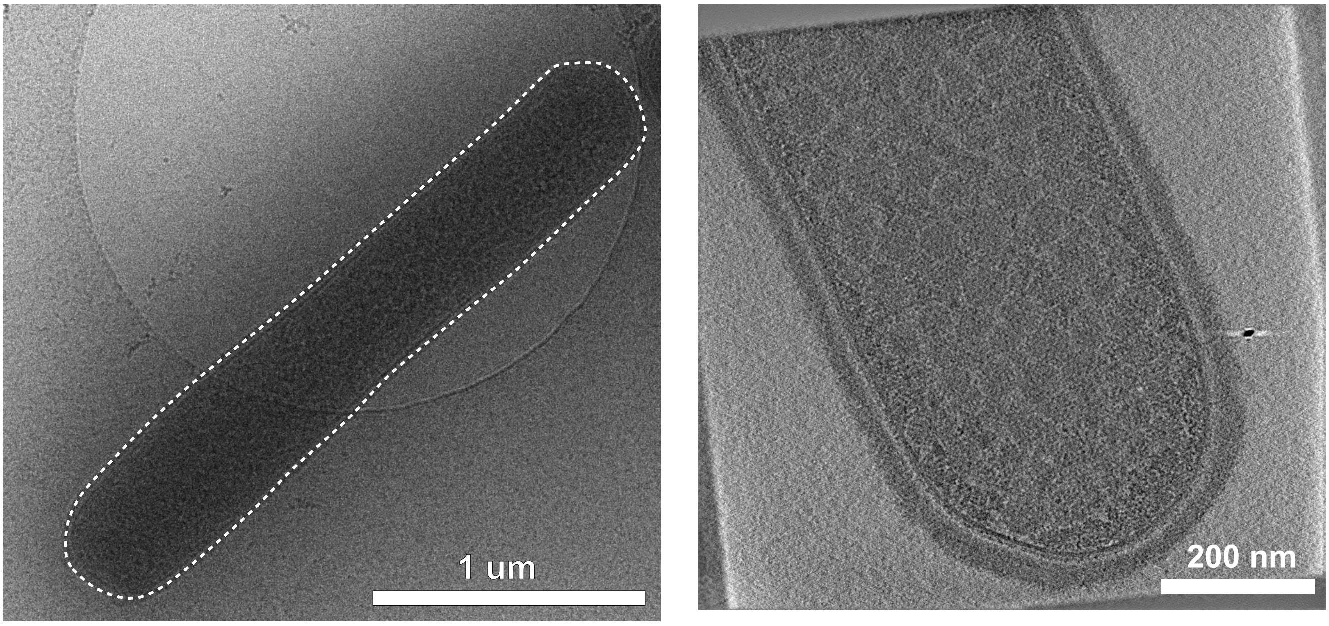

Streptococcus bovis ATCC 33317

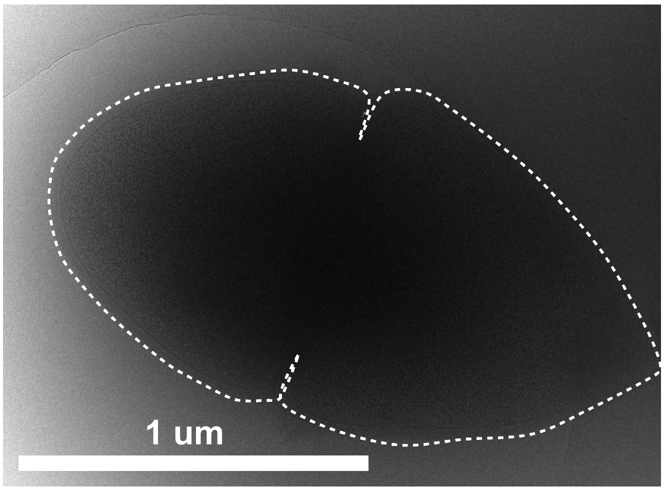


Streptococcus henryi A4

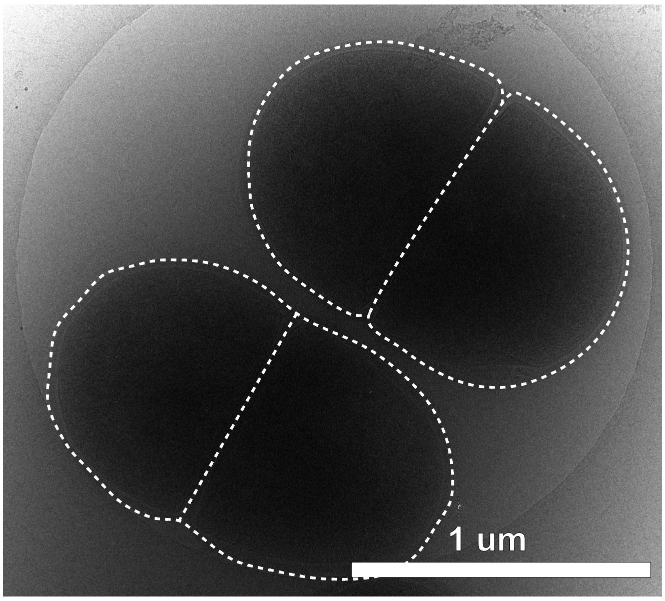


Succinimonas amylolytica DSM 2873

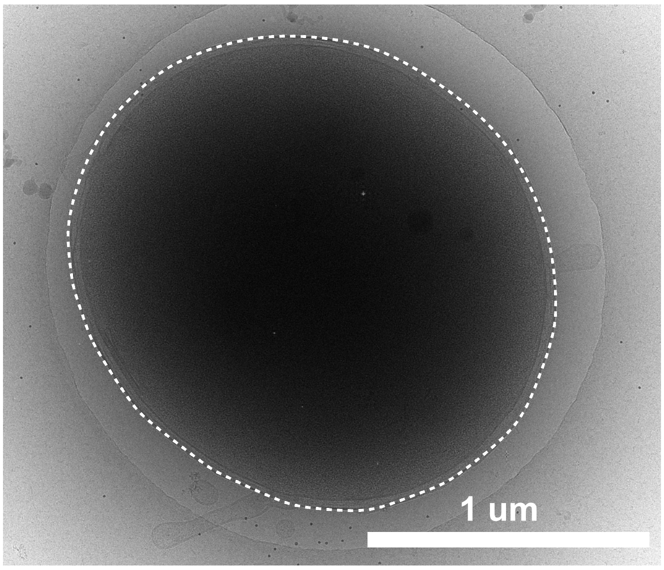


Succinivibrio dextrinosolvens DSM 3072

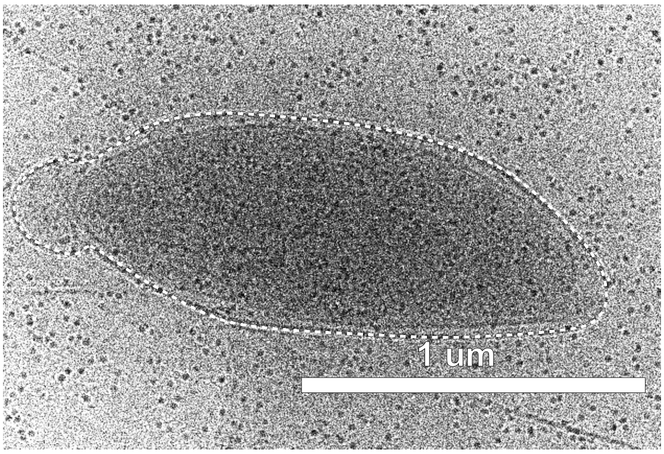


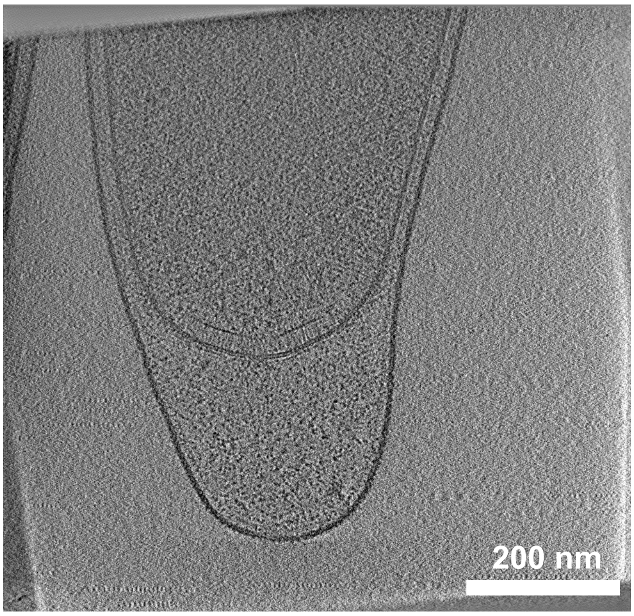

*Terrisporobacter glycolycus* KPPR-9

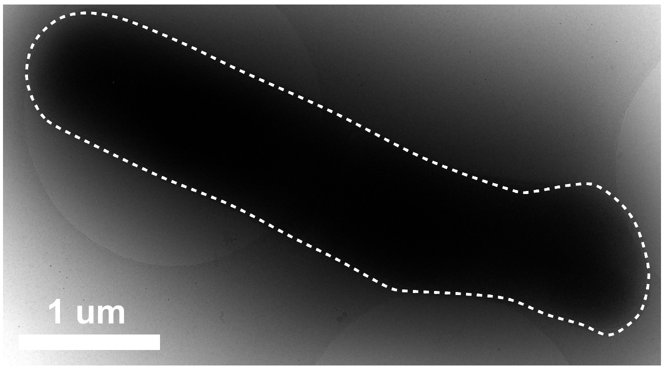


*Wolinella sp. / Vibrio succinogenes* ATCC 33567

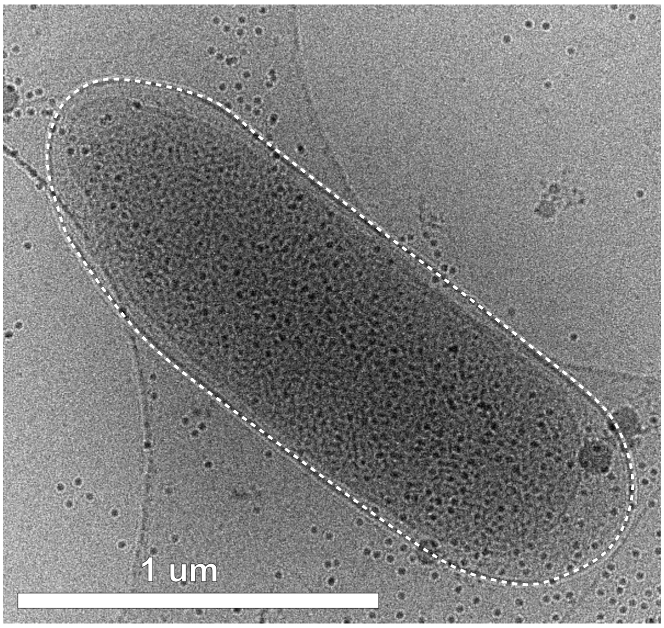


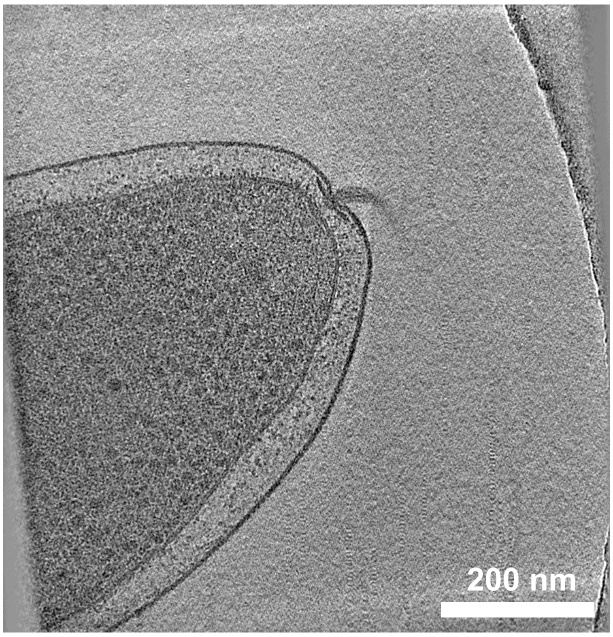
